# Highly dynamic mechanical transitions in embryonic cell populations during *Drosophila* gastrulation

**DOI:** 10.1101/2024.08.29.610383

**Authors:** Juan Manuel Gomez, Carlo Bevilacqua, Abhisha Thayambath, Maria Leptin, Julio M Belmonte, Robert Prevedel

## Abstract

During animal development, the acquisition of three-dimensional morphology is a direct consequence of the dynamic interaction between cellular forces and the mechanical properties of cells and their environment. While the generation and transmission of cellular forces has been widely explored, less is known about the dynamic changes in cell mechanical properties during morphogenesis. Here, we characterise and spatially map in three dimensions the dynamics of cell mechanical properties during *Drosophila* gastrulation utilising line-scan Brillouin microscopy. We find that cells in the embryo undergo rapid and spatially varying changes in their mechanical properties and that these differ in cell populations with different fates and behaviours. We identify microtubules as potential effectors of cell mechanics in this system, and corroborate our experimental findings with a physical model that underscores the role of localised and dynamic changes in mechanical properties to facilitate tissue folding. Our work provides the first spatio-temporal description of the evolving mechanical properties of cell populations during morphogenesis, and highlights the potential of Brillouin microscopy in studying the dynamic changes in cell shape behaviours and cell mechanical properties simultaneously in different cell populations in an intact organism.

## Introduction

A critical mechanism driving the three-dimensional acquisition of shape and form across the scales of living matter is the regulation of cell shape. This regulation is a direct consequence of the balance of forces produced by and acting on it, as well as the cell’s compliance to deformation, which in turn is ultimately determined by its mechanical properties [1, 2]. While the molecular mechanisms underlying the generation and the transmission of forces have been extensively studied [3–7], the dynamics of cell mechanical properties determined during the development of multicellular organisms are less understood. This is largely due to the challenges associated with measuring mechanical properties with high spatial and 3D resolution inside dynamic tissues.

The first major, embryo-scale morphogenetic event in metazoans is gastrulation, which connects cell fate specification with the coordinated acquisition of particular cellular cell shape behaviours [8]. In *Drosophila*, gastrulation begins with the formation of the ventral furrow (VFF), when ventral cells (mesoderm) accumulate medial-apical actomyosin, causing the apical constriction, shape changes, tissue folding and invagination of the mesoderm in an autonomous manner [3, 9, 10]. Once internalised, mesoderm cells undergo epithelial-mesenchymal transition (EMT) [11, 12](Supplementary Figure 1a,b and Supplementary Video 1). The remaining dorso-ventral (DV) cell populations respond differently to the internalisation of the mesoderm. Lateral cells (neuroectoderm) move *in-bloc* towards the ventral midline with minimal changes in their apical cellular geometry [8] (Supplementary Figure 1a,b and Supplementary Video 1). By contrast, dorsal cells, which will generate the embryonic dorsal ectoderm and the extra-embryonic amnioserosa, become squamous [8] (Supplementary Figure 1a,b and Supplementary Video 2). These differential cell shape behaviours during gastrulation suggest variations in mechanical properties among DV cell populations. Hence, *Drosophila* gastrulation constitutes an excellent model to study the intricate connection between mechanical dynamics and the acquisition of specific cell shape behaviours.

A range of experimental methods, often combined with theoretical models or simulations have been used to study the mechanical properties of cells in the early *Drosophila* embryo [8, 13–15]. Manipulations with ferrofluids and magnetic beads in cellularising embryos revealed differential mechanical properties of the apical cortex and the cytoplasm [14]. Apical tension fields were probed using laser microdissection along DV cell populations before and upon initiation of gastrulation, suggesting that differential mechanics emerge on the apical cortex of DV cell populations [8]. Although these studies advanced our mechanical understanding of early embryogenesis, they could not provide specific information on the dynamic changes of cell mechanical properties.

Recent progress in Brillouin microscopy, has enabled applications to living systems to assess cell mechanical properties in longitudinal and volumetric fashion with high spatial and temporal resolution [16]. Brillouin microscopy exploits the inelastic interaction between light and biological matter that causes a shift in photon energy (’Brillouin shift’) when photons are scattered by intrinsic collective acoustic vibrations within the sample. The shift in energy of the Brillouin-scattered light is related to the sound velocity of the probed material. From this, the so-called longitudinal modulus, which is a direct measure of elasticity of the biological material, can be calculated if the refractive index and mass density are known. However, even in the absence of these parameters, the Brillouin spectrum can be used as a proxy of visco-elastic properties [17]. Brillouin microscopy has been successfully applied to a variety of cell-biological questions, such as the investigation of intracellular biomechanics in whole living cells [18–20], the analysis of liquid-to-solid phase transitions in individual sub-cellular structures [19, 21], as well as the biomechanical assessment of entire tissues in living organisms [16, 17, 22].

Historically, Brillouin microscopy has been limited in its applications owing to the weak scattering interaction, which prevented the study of mechanical dynamics at the spatial and temporal scales relevant for morphogenesis (seconds to minutes over hundreds of micrometers). Recently, we introduced a line-scan approach to Brillouin microscopy that substantially improved its temporal resolution by two orders of magnitude and decreased significantly phototoxicity, thereby enabling the measurement of dynamics of mechanical properties of comparatively large tissue volumes within biologically-relevant timeframes for fast processes [16].

Here, we used Line-Scan Brillouin Microscopy (LSBM) to characterise and spatially map the dynamics of mechanical properties of the embryonic cell populations engaged in *Drosophila* gastrulation. We find differences in mechanical dynamics between the mesodermal and the ectodermal cell populations. In particular, the mesoderm undergoes biphasic mechanical dynamics, with an initial transient increase in Brillouin shift, followed by a softening during EMT. The observed increase in Brillouin shift was strongest in the sub-apical compartment of central mesodermal cells, where sub-apical microtubules become aligned, as confirmed by 3D-SIM super-resolution and live-cell confocal imaging. De-polymerization of the microtubules reduced the Brillouin shift during ventral furrow formation, suggesting that microtubules (MTs) are required for determining the mechanical properties of central mesodermal cells. To corroborate our experimental findings, we further built a physical model of VFF which confirmed the importance of sub-apical longitudinal stiffness to gastrulation and showed that the coordination of dynamic changes in central and peripheral cell stiffness improves furrow formation. In summary, we provide the first comprehensive description of evolving mechanical properties of embryonic cell populations during an embryo-scale morphogenetic event, and shed light onto the cellular components that may drive the observed fast and dynamic changes of cell mechanical properties during tissue folding.

## Results

### DV cell populations display differential mechanical dynamics during gastrulation

Cells along the DV axis change their shape in different ways (stage 6 [23]; Supplementary Fig. 1a-c, Supplementary Video 1,2), and we sought to study their mechanical properties using LSBM. We first imaged and analysed the dynamic changes of cell mechanical properties as indicated by the Brillouin shift within the mesoderm (ventral, Fig. 1a) focusing on a 16-cell wide field of cells along the ventral midline (8 cells on each side of the ventral midline; see Methods; Supplementary Figure 1a, Supplementary Video 1). We measured the Brillouin shift from the onset of VFF (stage 5b [23]) to the initial phase of epithelial-mesenchymal transition (EMT) (stage 8b [23]).

**Figure 1.**
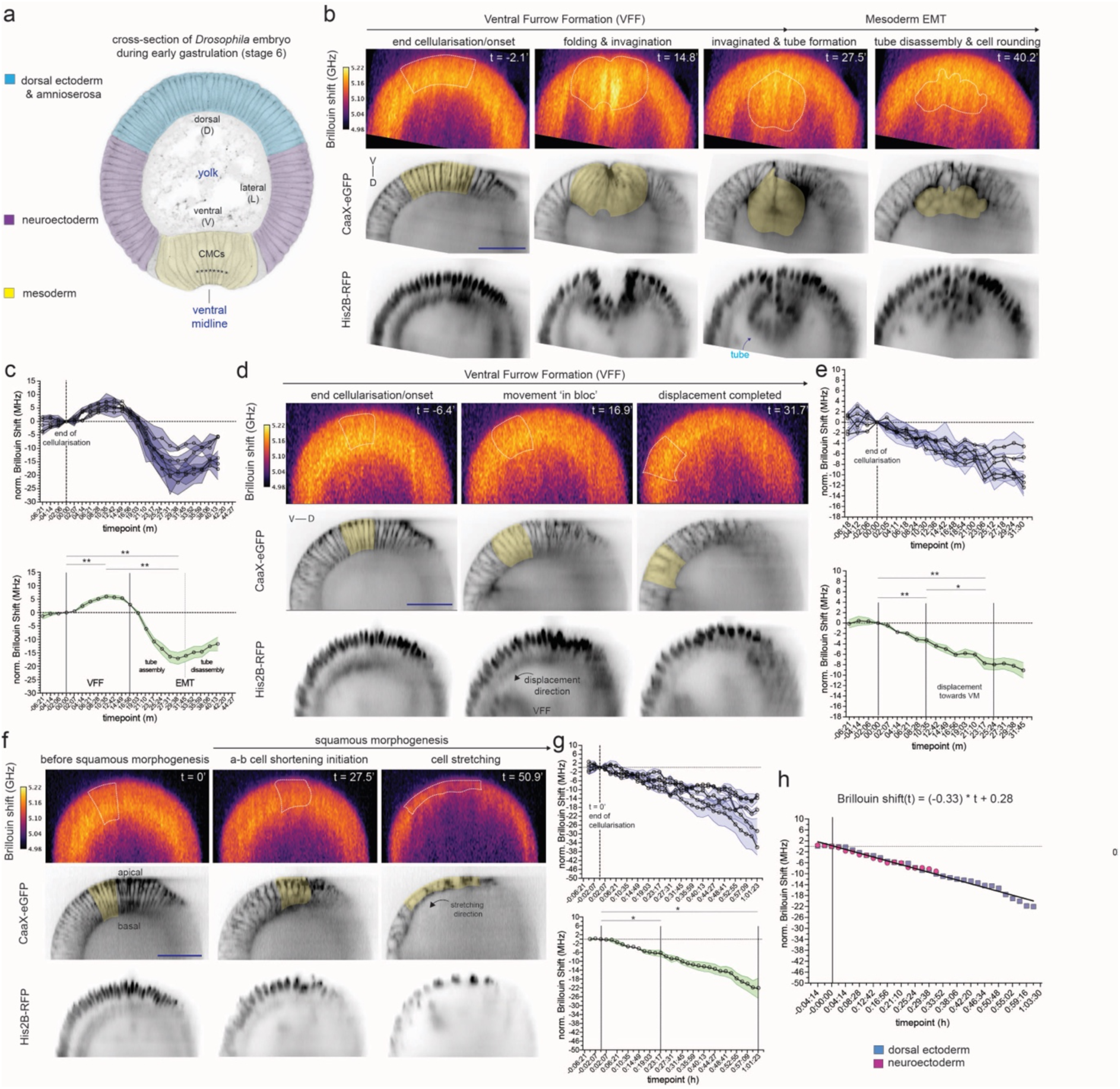
The dynamics of mechanical properties in dorso-ventral cell populations during gastrulation. **a.** Physical cross-section of a fixed embryo during the initiation of gastrulation (stage 6: VFF), indicating the position of the ventral midline (VM) and larger dorso-ventral cell populations. Asterisks (*) mark 8 cells within the central mesodermal. **b.** Brillouin shift (BS) maps of the mesoderm that correspond to particular stages of VFF (left and centre-left column) and epithelial-mesenchymal transition (EMT, centre-right and right column), obtained from average projections of three consecutive slices. Region shaded in yellow in CAAX-eGFP and enclosed with white dashed lines in the BS maps corresponds to the quantified areas used for assembling panel c. **c.** Quantification of the BS within the mesoderm during VFF and EMT. The BS was quantified within an initial 16-cell domain (8 cells on each side of the ventral midline, see Methods) as shown in panel b. Top panel: mean was obtained by quantifying the region of interest (initial 16-cell domain) in three YZ re-sliced sections (raw volume was re-sliced). BS was normalised to the onset of VFF. **d.** BS maps of the neuroectoderm that correspond to particular stages of neuroectoderm displacement towards the VM, obtained from averaging projections of two consecutive slices. Regions shaded in yellow in CAAX-eGFP and enclosed with white dashed lines in the BS maps correspond to the quantified areas used to assemble panel e. **e.** Quantification of the BS within the neuroectoderm during gastrulation. The BS was quantified within a 6-cell domain (see Methods) as shown in panel d. Top panel: mean was obtained by quantifying the region of interest (6-cell domain) in three YZ re-sliced sections (raw volume was re-sliced). BS was normalised to the end of cellularisation. **f.** BS maps of the dorsal ectoderm that correspond to the progress of squamous morphogenesis, obtained from average projections of two consecutive slices. Regions shaded in yellow in CAAX-eGFP and enclosed with white dashed lines in the BS maps correspond to the quantified areas used to assemble panel g. **g.** Quantification of the BS within the dorsal ectoderm during gastrulation. The BS was quantified within an initial 6-cell domain (see Methods) as shown in panel f. Top panel: mean was obtained by quantifying the region of interest (6-cell domain) in three YZ re-sliced sections (raw volume was re-sliced). BS was normalised to the end of cellularisation. **h.** Comparison of BS dynamics between the two subpopulations of the ectoderm: neuroectoderm (lateral; panels d,e) and dorsal ectoderm (dorsal; panels f,g) during gastrulation. Both the ectoderm and the neuroectoderm fit the same linear function. BS maps and BS dynamics shown in panels b-h were conducted in transgenic embryos with fluorescently labelled membranes (CAAX-eGFP: inverted grayscale, middle panel) and nuclei (His2B-RFP: inverted grayscale, bottom panel). Dorsal-(D)ventral(V) axis is marked in panels b, d and f to illustrate the position of the imaged embryo. Quantifications in panels c,e,g show: top panels: mean and SEM of the Brillouin shift in each of five quantified embryos; bottom panels: mean and SEM of the BS obtained from the 5 embryos shown in the top panel. SEM is the standard error of the mean. Scale bars are 50 μm. For statistical comparisons: * is p < 0.05, ** is p < 0.01.

When gastrulation starts we detected a transient increase in the Brillouin shift that peaks at the initiation of mesoderm invagination (Fig. 1a,b; t = 10:35 min, Supplementary Video 3; RM one-way ANOVA followed by multiple comparisons –FDR corrected-: p = 0.002), consistent with our own previous results [16]. We note that to convert the Brillouin shift to an absolute elastic modulus, the ratio between refractive index (*RI*) squared and mass density (*d*) is required. Measuring these parameters independently is difficult in a scattering and thick sample such as the *Drosophila* embryo. However, their ratio only varies a few percent for a wide range of proteins, nucleic acids and sugars [17] and therefore the Brillouin shift can be considered proportional to the longitudinal modulus. The most notable exception from the above assumption are lipid rich compartments [17]. Therefore, one possible cause for the observed increase in Brillouin shift within the mesoderm might be lipid droplets. To test this, we studied their distribution in mesodermal cells with a fluorescent marker for lipid droplets (YFP protein-trap in Lsd-2 [24, 25]). We found lipid droplets along the basal compartment of mesoderm cells during late cellularisation (Supplementary Fig. 1d, Supplementary Video 4) and the initiation of VFF, consistent with earlier observations in EM sections [26] and Raman loss microscopy [27]. Since the highest Brillouin shift was in a different region of the cell, this excludes the presence of lipids as the only explanation for the increased shift (Supplementary Fig. 1d, Supplementary Video 4), thus justifying the use of the Brillouin shift as a proxy for the longitudinal modulus in absence of spatial *RI* and/or *d* knowledge.

We then analysed the dynamics of mechanical properties in the mesoderm beyond VFF. Once VFF is completed, the invaginated mesoderm transiently arranges in a circular manner that resembles a tube (Supplementary Figure 1a, Supplementary Video 1) that spans across most of the anterior-posterior axis. When cells initiate the EMT, the tube disassembles, concomitant with the loss of a columnar shape and the acquisition of a spherical geometry (Supplementary Figure 1a, Supplementary Video 1). After the Brillouin shift peaks in the mesoderm (Fig. 1c; Supplementary Video 3), it decreases, concomitant with the completion of mesoderm invagination, and the initiation of EMT (Fig. 1b,c; Supplementary Video 3). During this phase cells lose their columnar shape and become round [11] (Fig. 1b; Supplementary Fig. 1a,b). When tube disassembly begins (Supplementary Fig. 1a), and before cells acquire a round shape (t = ∼31:45 min) mesodermal cells reach a maximum softening, as indicated by 17 MHz reduction in the Brillouin shift from the initiation of gastrulation (RM one-way ANOVA followed by multiple comparisons –FDR corrected-: p = 0.0014) and a 23 MHz reduction in the Brillouin shift (RM one-way ANOVA followed by multiple comparisons –FDR corrected-: p = 0.0007) from the initiation of mesoderm invagination, respectively (Fig. 1c).

Next, we analysed the Brillouin dynamics within a group of six cells in each of the ectodermal cell populations, the neuroectoderm and dorsal ectoderm. Although these two populations look very similar during the initial phase of VFF (Supplementary Fig. 1a, Supplementary Video 1), the dorsal ectoderm cells, but not the neuroectodermal cells, stretch in the direction of the ventral midline throughout stages 7-8 (Supplementary Video 2). From the end of cellularisation, both the neuroectoderm and dorsal ectoderm cells showed a linear reduction in the Brillouin shift (Fig. 1 d-g, Supplementary Videos 5,6). When neuroectoderm cells have completed their ventral displacement (i.e. end of stage 6), we measured on average a 7.7 MHz reduction in the Brillouin shift (t = 23:17 min; Fig. 1d,e and Supplementary Video 5; RM one-way ANOVA followed by multiple comparisons –FDR corrected-: p = 0.0027), which was similar to the reduction in Brillouin shift in dorsal ectoderm cells at the same time (−6.3 MHz, t = 23:17 min; Fig. 1e,g). Squamous morphogenesis continues beyond the period of the ventral displacement of the neuroectoderm. Thus, we followed the Brillouin shift further in time, and measured a reduction of 22 MHz in the dorsal ectoderm cells by the time it had become squamous (t = 01:01:23 hour; Fig. 1f,g and Supplementary Video 6; RM one-way ANOVA followed by multiple comparisons, FDR corrected: p = 0.0156). Finally, we tested if the mechanical dynamics of the neuroectoderm and dorsal ectoderm were the same (Fig. 1,e,g). Our analyses confirmed the Brillouin shift decreases at the same rate within both ectodermal cell populations during gastrulation (linear regression fit: Brillouin Shift (in MHz) = –0.33 * t + 0,28; stats dif. slope: p = 0.084; stats dif. intercept: p = 0.795).

### Brillouin shift dynamics in individual cells

Next, we focused on the association between changes in cell shape and the Brillouin shift at the level of individual cells. We again started our analyses with the 16 ventral mesodermal cell rows (Fig. 2a, top panel) in the early phase of VFF. At the end of cellularisation, the Brillouin shift profile was homogeneous (t = –2:07 min, Fig. 2a). When VFF started, the Brillouin shift became heterogenous (t = 2:07 min, Fig 2a). As VFF progressed, the Brillouin shift decreased slightly in peripheral mesodermal cells (positions |6-8|, Fig. 2a), but increased in cells closer to the ventral midline, i.e., central mesodermal cells, with a peak in the 3 cell rows located closest to the ventral midline (∼15 MHz increase, positions |1-3|, Fig. 2a). Central and peripheral mesodermal cells undergo different cell shape changes [6, 9]. Central mesodermal cells contract their apical sides, and experienced the largest increase in the Brillouin shift (Fig. 1a, Supplementary Video 1, Fig 2a –blue arrows-). Peripheral mesodermal cells stretch in the direction of the furrow [28] (Fig. 1a, Supplementary Video 1, Fig 2a –red arrows-) and showed a reduction in the Brillouin shift. Thus, the Brillouin shift profile in the mesoderm during the early phase of VFF correlates with cell behaviours.

**Figure 2.**
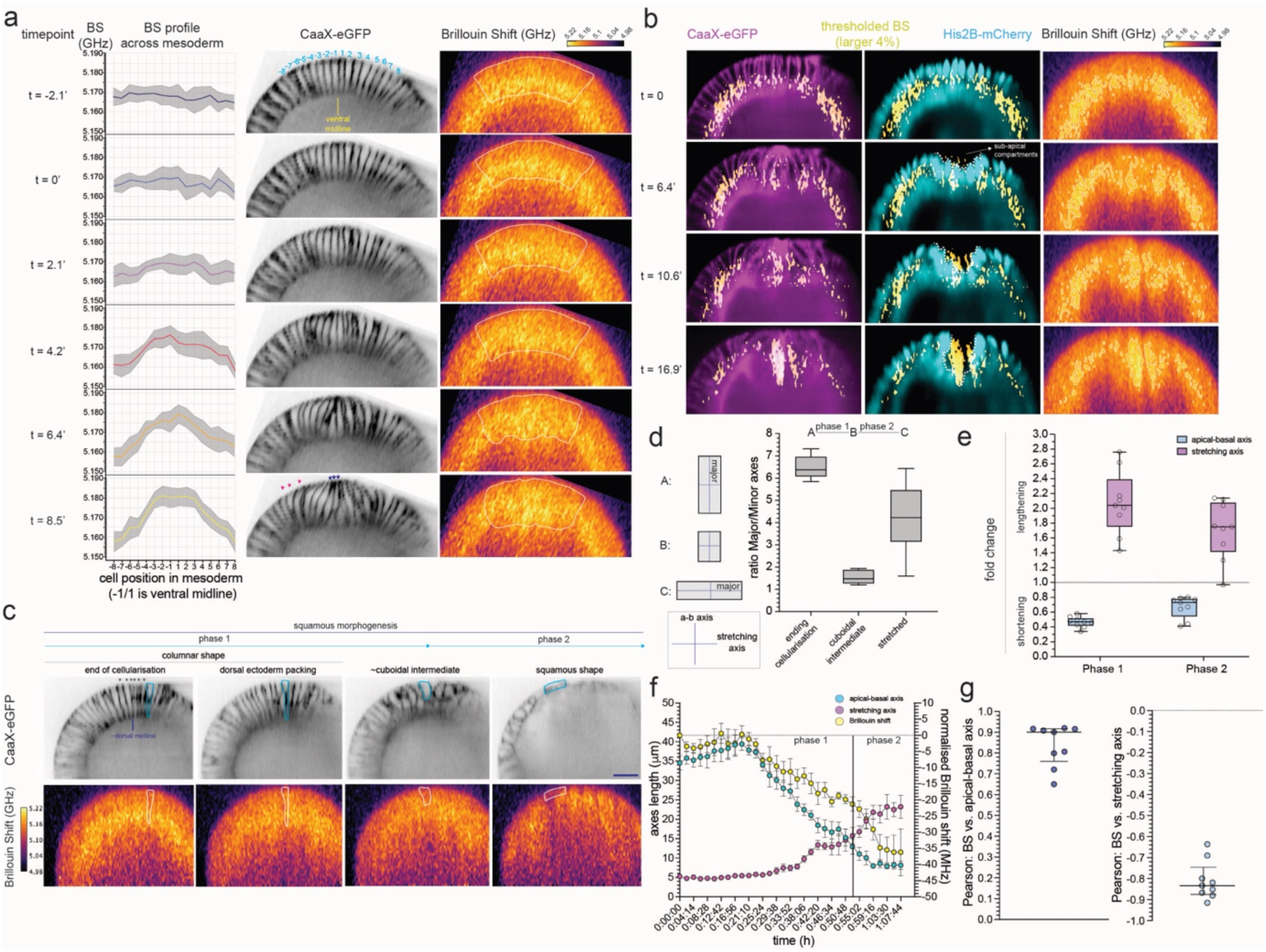
The dynamics of the Brillouin shift at the cellular level. **a.** Brillouin shift (BS in GHz) profile in the mesoderm during the early events of VFF (onset to folding initiation) in transgenic embryos with labelled membranes (CaaX-eGFP, centre panel: inverted grayscale). The profile was assembled by quantifying cell-by-cell the BS along a 16-cell field within the mesoderm (8 cells on each side of the ventral midline/VM). Cell position with respect to the VM is indicated by numbers (top panel: blue) that correspond to the cell numbers on the x-axis of the BS shift profile (left panel). Area enclosed with white dashed line is the quantified mesoderm field in the BS shift map (right panel). A line marks the mean BS shift for each cell position, and grey area Indicates the standard error of the mean (SEM). Three slices in each of four embryos were quantified. **b.** Subcellular localisation of largest values of the BS (largest 4% of the dynamic range, see Methods) in transgenic embryos with labelled membranes (CaaX-eGFP, left panel: magenta) and nuclei (H2B-RFP, centre panel: cyan). Filtered BS regions of the BS maps with values corresponding to the 4% largest were overlaid in CaaX-eGFP and H2B-RFP channels to visualise the localisation of the high shift. White dashed line in the centre panel shows the boundary between the nuclei and the sub-apical domain of central mesodermal cells (Fig. 1a, asterisks). **c.** Selected stages of squamous morphogenesis in dorsal ectoderm cells: columnar cells from the end of cellularisation (left) until the dorsal ectoderm becomes packed during the early phase of VFF (center-left), the formation of short columnar/cuboidal intermediate (center-right) and squamous cells (right). The boundary between phase 1 and phase 2 during squamous morphogenesis is set by the time point of the cuboidal intermediate. Cell boundary of an exemplary quantified cell (panels d-g) is shown in light blue (top panels) or white (bottom panels). Top panel shows membranes labelled with CaaX-eGFP, bottom panel shows BS map. Asteriks (*) mark cells considered to be amnioserosa cells (6 cells, 3 on each side of the estimated position of the dorsal midline) and thus, not included in our analyses. Scale bar is 25 μm. **d.** Box-whiskers plot showing the ratio between the major and minor cell axes in single dorsal ectoderm cells (N=9) at two time points: end of cellularisation (columnar shape), when the circularity value is maximal (largely cuboidal/isotropic shape) and the last quantified time point for each cell (squamous shape). Line within the box is the median, edges of the box 1^st^ and 3^rd^ quartiles and whiskers show maximum and minimum values for each cell. **e.** Box-whiskers plot showing the quantification of the fold change in apical-basal and stretching axes length from the end of cellularisation until cells reach the largest circularity value (’during phase 1’) and from the time point when cells reach the largest circularity value until they have become squamous (’during phase 2’). The apical-basal axis (a-b) is marked in cyan, the stretching axis is marked in magenta. Each dot is a quantified cell (N=9). **f.** Time course plot showing the cell-by-cell means of the Brillouin shift (yellow: normalisation to time = 0’, corresponding to the end of cellularisation), the apical-basal (a-b, cyan) and stretching (magenta) axis lengths (in μm) during squamous morphogenesis of the dorsal ectoderm. N=7, see Methods. Bars show standard error of the mean (SEM). **g.** Scatter plot showing the Pearson correlation values for each analysed cell (N=9) between the change in the median of the normalised Brillouin shift (in MHz) and the change in the length of the apical-basal (left, cyan) and stretching (right, magenta) axes during the time course of squamous morphogenesis. BS is Brillouin shift. For statistical comparisons * is p < 0.05, ** is p < 0.01.

We further explored the subcellular dynamics of the transient increase in the Brillouin shift in central mesodermal cells for which we filtered the pixels in the Brillouin shift maps that had values in the top 4^th^ percentile and analysed their subcellular distribution. When mesoderm folding began, these filtered Brillouin shift pixels were detected between the apical membrane and the apical side of the nucleus, a compartment to which we will refer as ‘sub-apical’ (Fig. 2b, white dashed line, Supplementary Video 7). When folding progressed to invagination (t = 16.9’; 16:56 min) this entire area of the cell accumulated the pixels with highest Brillouin shift values.

Next, we analysed whether the reduction in Brillouin shift during the squamous morphogenesis of the dorsal ectoderm correlates with changes in cellular geometry. We divided the transition from a columnar to squamous shape in two phases (Fig. 2c,d and Supplementary Fig. 2a,b). In the initial phase (phase 1) the columnar cells shortened in the apical-basal axis and expanded in the stretching axis (Fig. 2d,e), generating roughly isotropic cells (Fig. 2d). This was followed by a second phase (phase 2), in which cells stretched along the DV axis and shortened further on the apical-basal axis until cells acquired a squamous shape (Fig. 2d,e).

We started to detect changes in the Brillouin shift soon after these cells have changed the lengths of the apical-basal axis and the stretching axes (Fig. 2f, t = 21:10 min). In each cell, we measured a continuous reduction in the median Brillouin shift that correlates positively with the reduction in apical-basal axis length (Fig. 2g, left panel), and inversely with the stretching of the cell along the DV axis (Fig. 2g, right panel). In summary, we found that the changes in cell shape correlate with changes in the Brillouin shift, and thus, in the mechanical properties of those cells.

### The distribution of actomyosin and actin binding proteins does not correlate with the high Brillouin shift measured during VFF

Changes in elasticity must be mediated by cellular components, and an important system in cell mechanics is the actomyosin network. The Brillouin shift profile (Fig. 2a) resembled the gradient of activated myosin within the mesoderm [6]. Mesodermal cells located closer to the ventral midline have higher levels of activated apical actomyosin, and thus, contractility, decreases in a gradient from central to peripheral cells [6]. Therefore, our results point to the cytoskeleton as the cellular component responsible for the measured mechanical properties during VFF and in particular, suggest actin as a possible contributor to the transient increase in the Brillouin shift within central mesodermal cells.

To explore an involvement of the actin cytoskeleton on mechanical dynamics in the mesoderm, we first looked at the colocalisation of actomyosin and the Brillouin shift in embryos carrying fluorescently labelled versions of myosin (sqh-mCherry) and F-actin (UtrophinABD-GFP). When the mesoderm folds (t = 10’), non-muscle myosin and F-actin accumulate on the apical side of cells [3] (Fig. 3a, arrows, Supplementary Video 8). A small fraction of the high Brillouin shift colocalised with actomyosin during the invagination of the mesoderm (t = 11:40 – 15:00 min, Fig. 3a, Supplementary Video 8). However, the transient increase in the Brillouin shift was detected largely basal to the apical cortex, i.e. beyond the area directly at the cortex where actomyosin is concentrated (Fig. 3a, white dashed area; Supplementary Video 8). This finding is consistent with the observation of a transient high Brillouin shift in myosin-independent folding events [29]. During dorsal fold formation (Fig. 3b, Supplementary Video 9), we also observed an increase in the Brillouin shift, thereby further supporting that actomyosin is not the sole cellular component responsible for the transient increase in Brillouin shift.

**Figure 3.**
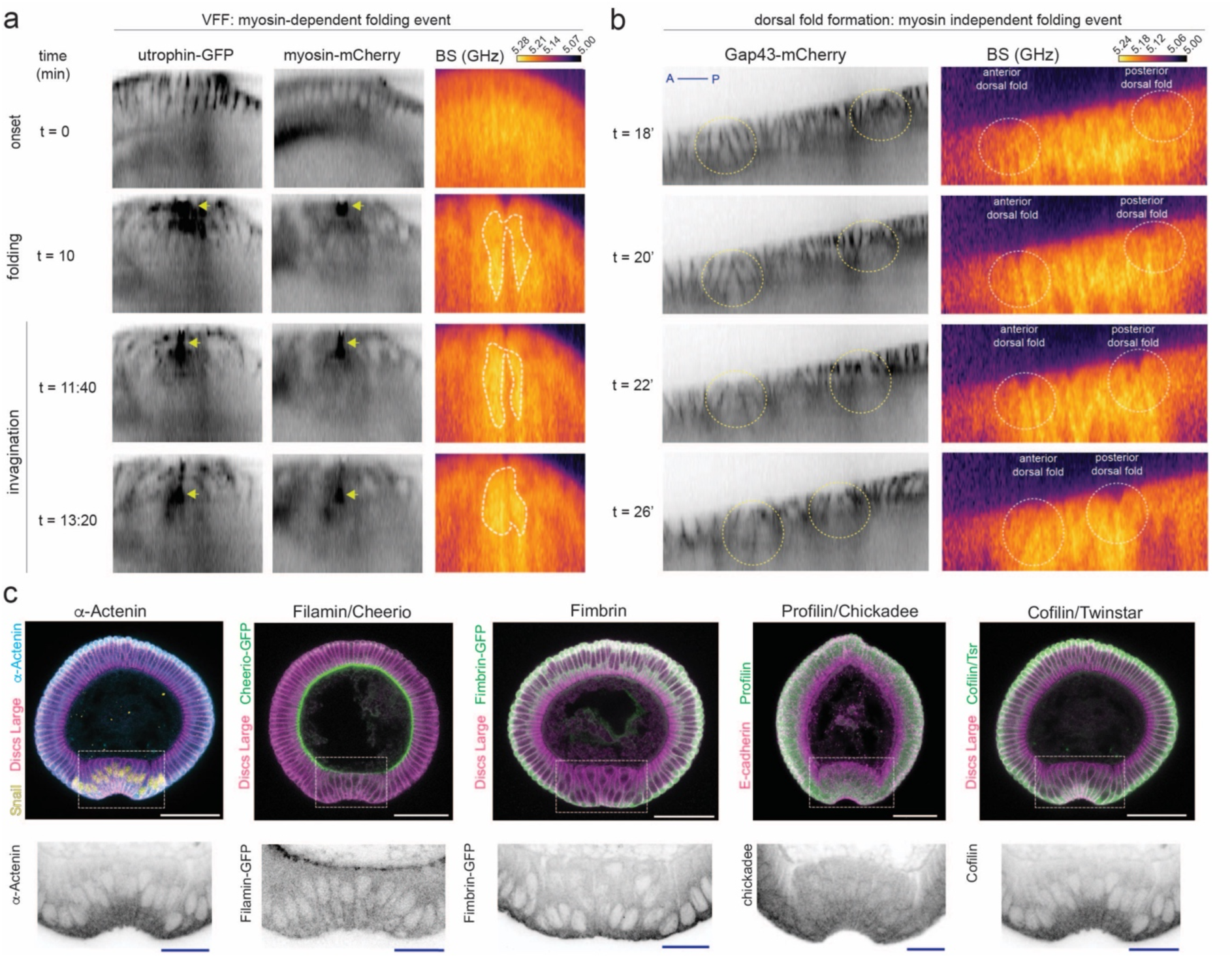
Comparison between actin and actin binding proteins distribution and transient increase in Brillouin shift. **a.** Comparison of the localisation of actomyosin and Brillouin shift (right panel) during VFF in transgenic embryos with labelled F-actin (utrophin-GFP, left panel: inverted grayscale) and myosin light chain (myosin-mCherry, centre panel: inverted grayscale). Frames show central mesodermal cells (Fig. 1a, asterisks) at three stages of VFF progression (onset, folding initiation and invagination). Arrows indicate active myosin light chain and F-actin accumulation along the apical side of cells. Area of the mesoderm circled with a white dashed line shows increase in the Brillouin shift during folding and invagination. **b.** BS shift maps (right panel) from mid-sagittal views of dorsal ectoderm cells (Fig. 1a, blue) during dorsal fold formation (DFF) in transgenic embryos with labelled membranes (gap43-mCherry, left panel: inverted grayscale). Dashed circles (left panel: yellow; right panel: white) indicate tissue areas in which dorsal folds are forming. Time is normalised to the end of cellularisation. Anterior (A) is left, posterior (P) is right. **c.** Visualisation of actin binding proteins (top panels: cyan: α-Actenin, green: Filamin/Cheerio, Fimbrin, Profilin/Chickadee and Cofilin/Twinstar; bottom panels: inverted grayscale) in physical cross-sections of fixed embryos undergoing VFF (folding phase). Membranes were stained with antibodies against Discs Large (magenta) except in Profilin/Chickadee stainings, which were co-stained with an antibody against E-Cadherin. Nuclei expressing Snail (yellow) mark mesodermal cells. White dashed squares indicate magnified regions that correspond to the folding mesoderm shown in insets. Scale bars are 50 μm for embryonic D-V cross-section and 20 μm for insets.

The dynamic polymerisation and crosslinking of actin filaments with each other regulates actomyosin function and thus, could be the source of the high Brillouin shift observed during VFF. To study this possibility, we analysed the distribution of the bundling actin-actin crosslinkers α-Actinin and Fimbrin, the mesh crosslinker Cheerio/Filamin, and the regulators of F-actin dynamics Profilin/Chickadee and Cofilin/Twinstar within the mesoderm (Fig. 3c, Snail-positive cells) in physical cross-sections of fixed embryos undergoing VFF. These actin binding proteins were found in different subcellular localisations. α-Actinin, Fimbrin, Chickadee/Profilin and Cofilin/Twinstar were present throughout the cytoplasm (Fig. 3c). In contrast, Cheerio/Filamin was enriched in the apical and apical-lateral cortex and at the cellularising front of blastoderm cells (Fig. 3c). Additionally, Cheerio/Filamin was also differentially distributed along the DV axis (Fig. 3c), suggesting Cheerio/Filamin has different functions in the different cells along the DV axis. None of these actin binding proteins were specifically enriched within the subapical compartment of central mesoderm cells (Fig. 3c). These results suggest that neither crosslinked actin nor actomyosin are directly responsible for the transient stiffening of central mesoderm cells.

### The role of microtubules in the determination of mechanical properties during VFF

The other major system involved in cell mechanics is that of the microtubules. Networks of MT binding proteins and tubulin subunits are regulated through phosphorylation during gastrulation [28], and MTs are involved in controlling nuclear localisation and shape homeostasis in the context of tissue folding [28, 30, 31].

We looked at the organisation of MTs in physical cross-sections of fixed embryos before and during VFF. Before gastrulation, blastoderm cells have a sub-apical and a basolateral population that differ in their organisations[28, 32] (Fig. 4a). When apical constriction starts, the nuclei move basally [28] together with the centrosomes (labelled with the centriole component Asterless [33]; Fig. 4a, arrows). The movement of the centrosomes and the nuclei towards the basal side of cells enlarged the subapical compartment in central mesodermal cells (Fig. 4a, insets). During VFF, this enlarged sub-apical compartment became filled with long MTs that aligned with the cellular apical-basal axis (Fig. 4b).

**Figure 4.**
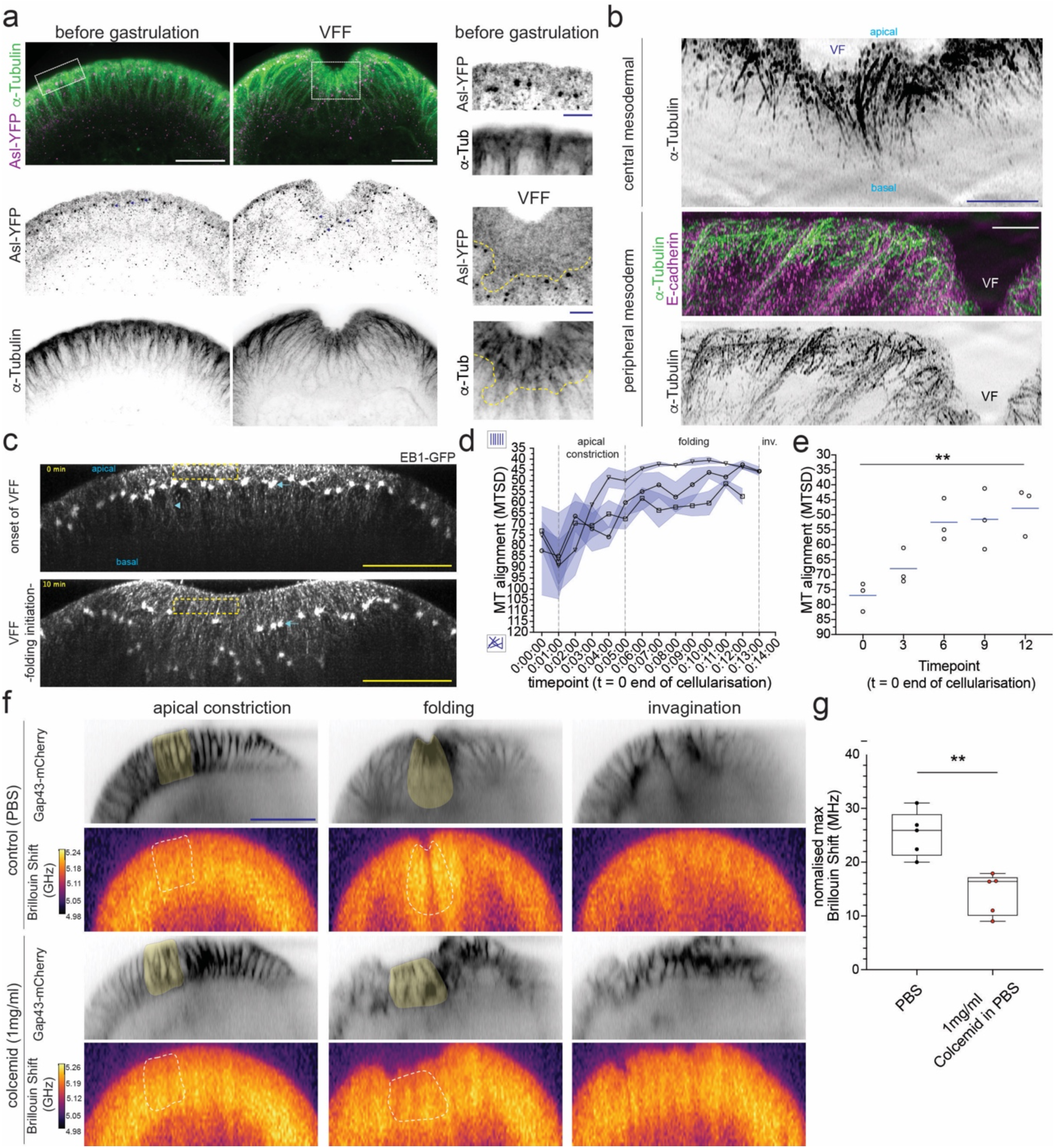
A role for microtubules in the high Brillouin shift within central mesodermal cells. **a.** Visualisation of microtubules (α-Tubulin, top panel: green; bottom panel: inverted grayscale) and centrosomes (Asterless-YFP, top panel: magenta; middle panel: inverted grayscale). Images correspond to max-projections of 10 consecutive slices of physical cross-sections of fixed embryos before and during gastrulation (VFF-folding phase). White dashed squares indicate magnified regions in insets (right panel), yellow dashed line illustrates an approximate boundary for the sub-apical compartment based on asterless larger puncta (centrosomes). Scale bars are 25 μm and 10 μm for insets. **b.** Super resolution imaging (OMX 3D-SIM) of sub-apical microtubules (top panel: inverted grayscale; middle panel: green; bottom panel: inverted grayscale) in fixed embryos undergoing VFF. Top panel shows sub-apical microtubules in central mesodermal cells, middle and bottom panel shows sub-apical microtubules in peripheral mesodermal cells (stretching or ‘hinge’ cells). Images correspond to max-projections of 30 consecutive slices acquired. VF = ventral furrow. Scale bars are 5 μm. **c.** Spinning disk images (max-projected) of the ventral side from living embryos carrying an EB1-GFP (grayscale) transgene to label the growing plus ends of microtubules. Top panel shows the mesoderm area at the onset of gastrulation (t = 0’); bottom panel shows the mesoderm when ventral furrow starts to be formed (t – 10’). Dashed yellow rectangles mark the area of the cells that was used for the quantification of the degree of alignment of microtubules (panel d,e). Green arrows: centrosomes before (top) and during VFF (bottom). Blue arrow: position of the ventral midline (VM). Blue arrowhead: basal-lateral microtubules. Note basal-lateral, but not sub-apica microtubules are aligned at the onset of VFF. Ventral is top. Scale bars are 25 μm. **d.** Quantification of microtubule alignment within the sub-apical compartment (yellow dashed rectangle). MTSD was calculated from the onset of VFF (t = 0’, end of cellularisation) until the initiation of mesoderm invagination (t = 13’-14’). The MTSD parameter decreases when MTs become aligned. Each line is one embryo (an independent experiment), with an average and standard error of the mean (SEM) from 5 max-projections, each projection derived from 20 YZ re-slices. **e.** Scatter-plot of selected timepoints from panel d: (0’, 3’, 6’, 9’ and 12’), showing the mean MTSD for each embryo (dots) and the median across the three embryos (blue line). The statistical differences in MTSD median across these selected timepoints were evaluated with a non-parametric one-way ANOVA (Friedman) test, followed by multiple comparisons (FDR corrected). ** is p < 0.01. **f.** BS maps at apical constriction (left panel), mesoderm folding (centre panel) and invagination (right panel) in control (PBS, top panel) and Colcemid-treated embryos (Colcemid 1mg/ml in PBS, bottom panel) in embryos with labelled membranes (Gap43-mCherry, inverted grayscale). Region shaded in yellow in Gap43-mCherry channel and enclosed with white dashed line in the BS maps corresponds to the quantified areas used for assembling panel d. Scale bar is 50 μm. **g.** Box-whiskers plot showing the quantification of the maximum BS using median, 1^st^ and 3^rd^ quartile, measured in control and Colcemid-treated embryos from the onset of VFF to 35’ after the end of cellularisation. The BS was quantified within an initial 6-cell domain of central mesodermal cells centered at the ventral midline (Fig. 1a). Each dot represents the mean of the 6-cell domain in a particular embryo (black dot is control, red dot is Colcemid-treated). Mean was obtained by quantifying the region of interest (6-cell domain) in a single slice in each of 5 embryos (5 independent experiments). BS was normalised to the end of cellularisation. Statistical comparison was performed using a Mann-Whitney test (non-parametric). For statistical comparisons: * is p < 0.05, ** is p < 0.01.

We also analysed the dynamics of sub-apical MTs in living embryos using spinning-disk confocal microscopy and a transgenic line that labels the MT plus-end tracking protein EB1 (EB1-GFP [34]) To quantify the degree of alignment of MTs with each other, we used a Microtubule Standard Deviation [35] (MTSD) which measures the variability of the EB1-GFP signal from its average direction. We found that the EB1-GFP signal became increasingly organised within the sub-apical compartment during VFF (Fig. 4c-e, Supplementary Video 11). Most of the reorganisation of the EB1-GFP signal occurred between the onset of VFF (t = 0’, MTSD = 77) and the initial phase of fold formation (t = 6’, MTSD = 53). During the later folding phase, sub-apical MTs further aligned with each other and the MTSD parameter reached a minimum when folding had progressed to the invagination of the mesoderm (t = 12’, MTSD = 48; Supplementary Video 11, Fig. 4d,e, one-way non-parametric ANOVA –Friedman test-followed by multiple comparisons –FDR corrected-: p = 0.0045). In summary MTs undergo a fast reorganisation during early VFF, filling the sub-apical compartment of central mesodermal cells with aligned and dynamic MTs. This coincides with the time when we measure the peak in Brillouin shift (Fig. 1b,c, Fig. 4d, Supplementary Video 3,11).

We reasoned that the reorganisation of sub-apical MTs parallel to the apical-basal axis and thus parallel to the probing direction of the Brillouin microscope, may be the origin of the measured high Brillouin shift during mesoderm invagination. To test this hypothesis, we depolymerised MTs by delivering Colcemid during mid-to-late cellularisation (see Methods). We focused on the six central mesodermal cell rows with the highest Brillouin shift (Fig. 2a). As previously reported, Colcemid treatment results in defects in cell shape and nuclear positioning, and ultimately the mesoderm fails to invaginate [28, 31] (Fig. 4c, Supplementary Video 10). During the initiation of VFF, the Brillouin shift within the six central mesodermal cell rows of Colcemid-treated embryos increased similarly to untreated embryos (Fig. 4f, Supplementary Fig. 2c). However, Colcemid-treated embryos reached a maximum Brillouin shift that was smaller than in control embryos (median: 16.4 MHz vs. 25.9 MHz, Mann-Whitney test p = 0.0079, Fig. 4f,g, Supplementary Video 10). Furthermore, the Brillouin shift in Colcemid-treated embryos stabilised close to its maximum; in control embryos the increase was transient (Supplementary Fig. 2c).

Our findings suggest that the apical constriction and folding of the mesoderm enables the stiffening of the central mesoderm, through the generation of an enlarged sub-apical compartment, ranging from the apical cortex to the centrosomes (Fig. 4a, Fig 4c –arrows-). This space could enable dynamic MTs to become highly aligned. To test the connection between the formation of the folding event itself and the high Brillouin shift, we analysed *twist* mutant embryos, which are known to impair but not fully abrogate apical constriction, and which engage a few mesodermal cells in fold formation [9] (Fig. 4e, Supplementary Video 12). Fold formation is delayed in *twist* mutants (t = 53.8’; 53:48 min) compared to control embryos (t = 20.7’; 20:42 min) and the transient increase in Brillouin shift is reduced (Supplementary Fig. 2d, Supplementary Video 12). Hence, our results suggest that engaging the whole mesoderm and achieving full contractility are required for the changes in the mechanical properties as measured during VFF.

### A model for the facilitation of tissue folding by tissue stiffening

Our observation of a transient increase in the Brillouin shift in the context of folding events, including VFF, dorsal fold formation (Fig. 3B), posterior midgut invagination [16] and neurulation [36], suggests that such a transient and spatially restricted increase in stiffness might be important for the proper progression of folding events. To test this hypothesis, we developed a physical toy model of *Drosophila* VFF within the Cellular Potts framework [37] in which we incorporated known biological qualities such as apical-medial actomyosin contractility, and spatially and temporally varying stiffness, as qualitatively informed by our measurements.

The model simulated the ventral half of the anterior-posterior cross-section of the embryo, encompassing a mesoderm width of 19 cells centred at the VM (Fig. 5a). Here, cells were divided into three compartments: apical, core and basal domains, reflecting the apical– basal polarity of blastoderm cells (Fig. 5b). Apical constriction was implemented using spring-like forces linking the apical sides of the cells (Fig 5b, first term), where the strength of the contractile force varies over time according to measured myosin levels within the mesoderm [6] (Fig. 5c). Assuming a linear longitudinal stress/strain relation, we use a similar approach to model the stiffness of the cells along their apical–basal axis (Fig 5b, second and third terms). The two *λ* parameters are proportional to the longitudinal modulus of the sub-apical and sub-basal regions of the modelled cells (see SI for a full model description).

**Figure 5.**
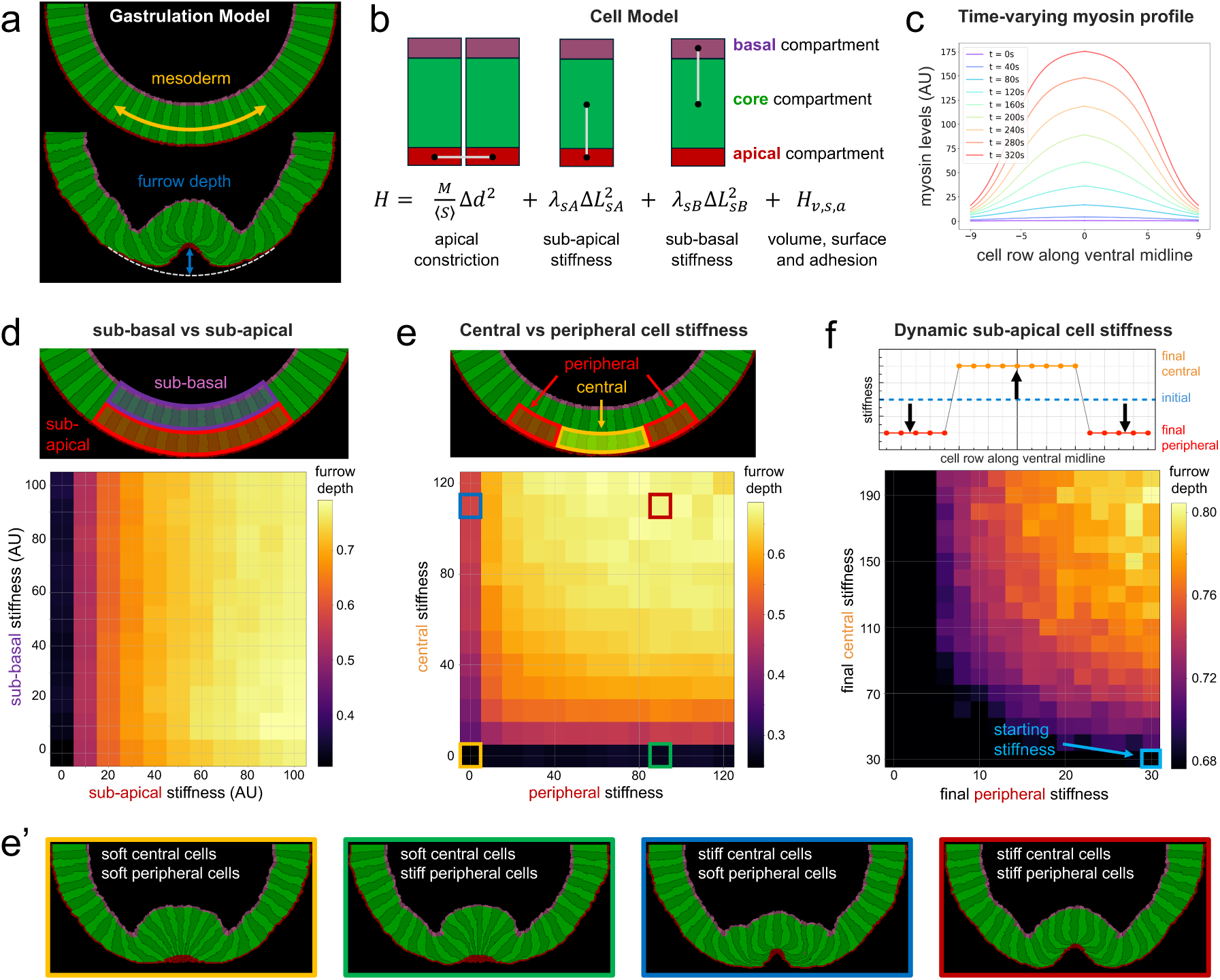
A physical model of *Drosophila* ventral furrow formation. **a.** *Top*: Initial condition of the 2D model with 47 cells representing the ventral half of the *Drosophila* embryo, including 19 mesodermal cells (yellow arrow range) and 28 neuroectodermal cells. *Bottom*: Schematic of the furrow depth metric, measured as the normalised distance between the apical-most coordinate of the central mesodermal cell and the lowest point of the mesoderm from the initial condition in units of initial cell height in the apical-basal axis. **b.** Each cell was modelled with 3 compartments: apical, core and basal. Hookean links connecting the centre of mass of neighbouring compartments are used to model apical constriction between adjacent mesodermal cells, and sub-apical and sub-basal longitudinal stiffness of mesodermal cells. Those constraints are part of an effective energy equation (*H*) that guides the model evolution and also include an adhesion, surface and volume terms as indicated. **c.** Experimentally measured [6] time varying spatial profiles of myosin concentration used to impose apical constriction on mesodermal cells. **d.** Furrow depth as a function of sub-apical and sub-basal longitudinal stiffness of mesodermal cells. Sub-basal stiffness variation has little effect on furrow formation, while higher values of sub-apical stiffness lead to deeper furrows. Each grid point value shows the average of 30 simulations. **e.** Furrow depth as a function of sub-apical longitudinal stiffness of central and peripheral cells. When either cell population is too soft, furrows fail to form (c.f. examples in panel **e’**). Higher values of central cell stiffness promote furrow formation even for relatively low values of peripheral cell stiffness. Sub-basal stiffness was the same for both cell populations. Each grid point value shows the average of 30 simulations. **e’.** Representative simulation snapshots of 4 cases from panel **e** (indicated by corresponding outline colour). **f.** Furrow depth as a function of increasing sub-apical central cell longitudinal stiffness and decreasing sub-apical peripheral cell stiffness. Grid coordinate indicates final values of central and peripheral cell stiffness. Colour bar cropped at the furrow depth value of simulations where stiffness of both cell populations remains constant at 30. For most points in the grid, final furrow depth was larger than the corresponding coordinates in panel **e** where the cells start the simulation with a higher(lower) central(peripheral) stiffness (see Supplementary Fig. 3d). Sub-basal stiffness was kept constant and the same for both cell populations. Each grid point value shows the average of 30 simulations.

We performed stepwise analyses of the effect of the previously identified variations in mechanical properties of cells during VFF: (i) a spatially restricted stiffness within each mesodermal cell, (ii) a stiffness gradient from central mesodermal to peripheral mesodermal cells and (iii) a dynamic change in cell stiffness in the order of minutes that recapitulates qualitatively our Brillouin measurements. We used the furrow depth (Fig. 5a) as a metric to quantify the extent of fold formation, and therefore as a proxy for proper fold progression. When comparing the effect of differential stiffness along the apical–basal axis of the mesodermal cells (Fig. 5d, top) we found that the sub-apical stiffness had a significantly larger effect on the furrow depth than sub-basal stiffness (Fig. 5d, bottom). Cells with a very small longitudinal stiffness were too soft and failed to form a furrow. Conversely, increased stiffness of sub-apical regions always favoured deeper furrow formation for the range of values that we tested (see Supplementary Fig. 3a).

To assess the role of the stiffness profile along the mesoderm (Fig. 2a), we fixed the sub-basal stiffness and set different sub-apical stiffnesses for the 9 central mesodermal cells and the 5 peripheral cells flanking the central cells on each side (Fig. 5e, top). When the central cells are very soft, furrow formation is blocked regardless of peripheral stiffness values (Fig. 5e’, first 2 panels; Supplementary Video 13 and 14). A high central cell stiffness with very soft peripheral cells promotes some furrow formation, but the high deformability of peripheral cells prevents a deeper furrow (Fig. 5e’, third panel; Supplementary Video 15). Optimal furrow formation occurred when both central and peripheral cells were sufficiently stiff with best results achieved when both stiffnesses were high (Fig. 5e’ last panel; Supplementary Video 16). From this we conclude that longitudinal cell stiffness is necessary to translate apical constriction movements into wedge cell shapes that lead to furrow formation.

Finally, we demonstrate the importance of dynamic changes in cell stiffness by varying the stiffness of the mesodermal cells over time to mimic the measured changes in stiffness during VFF (Fig. 2a; Fig. 5f, top panel). Starting from a constant value for all cells, we varied sub-apical stiffness over time to resemble the timing seen in the experiments, with peripheral cells decreasing in stiffness over time, followed by central cells increasing in stiffness over time (Supplementary Fig. 3c). Decreasing peripheral stiffness hinders furrow formation in most regimes and increasing central stiffness facilitates furrow formation (4% to 40% increase in furrow depth, depending on the starting sub-apical stiffness). The best results were with peripheral cells decreasing their stiffness up to 33% and the central cells increasing their stiffness above 200% from the starting value. Most strikingly, in most cases, dynamically changing cell stiffness over time produces deeper furrows than starting all the cells with a higher or lower central or peripheral stiffness, suggesting that dynamical changes of stiffness favour furrow formation (Supplementary Fig. 3d).

## Discussion

By utilising advanced line-scan Brillouin microscopy, we found that different cell populations within the *Drosophila* embryo exhibit rapid and spatially varying changes in mechanical properties during gastrulation, with divergent properties between the mesoderm and the whole ectoderm. Moreover, we observed a biphasic dynamic in mechanical properties in mesoderm cells, characterised by an initial stiffening followed by a softening during epithelial-mesenchymal transition (EMT). This dynamic is particularly pronounced in the sub-apical compartment of central mesodermal cells. Combining genetics, live-cell as well as super-resolution imaging, together with chemical perturbation, we were able to show that likely microtubules are the key determinants of the observed transient stiffening. Our physical model of ventral furrow formation corroborates these findings, underscoring the critical role of localised stiffness changes in facilitating tissue folding.

The timescales of the cellular and subcellular processes that regulate morphogenetic events, in particular those that are driven by cell shape changes, range from milliseconds to minutes. However, Brillouin microscopy measures the longitudinal modulus in the GHz regime, and thus, measures molecular mechanics that occur on nanosecond time-scales [38, 39]. While Brillouin spectroscopy was shown to be able to reveal both the elastic and viscous properties of biopolymers that are central to the structure and function of biological tissues [18, 40, 41], this difference in timescale still raises the question whether Brillouin microscopy is suitable for studying morphogenesis [17, 41]. Previous work has compared Brillouin microscopy with lower frequency techniques such as Atomic Force Microscopy (AFM), a technology that measures the Young’s modulus on time-scales more closely related to those of morphogenesis. While earlier work found positive correlations between the Brillouin frequency shift and the Young’s modulus in cells [18] and tissues [17, 41], recent evidence discourages a priori assumptions and advocates for the combined use of both methods for a more comprehensive mechanical characterisation of biological matter [42].

Our mechanical characterisation of cells along the dorsal-ventral axis fits well with measurements of mechanical properties using AFM in cells undergoing similar biological processes. The softening of cells undergoing EMT (Fig. 1) is consistent with the reduction in the elastic modulus that was observed in various models of cancer invasiveness and EMT [43–45]. The stiffening of contractile mesodermal cells during VFF is consistent with an increase in the elastic modulus in contractile cells, such as cardiac and skeletal muscle cells [46, 47], although the mechanisms driving stiffness might be different. Previous work suggested cells in the follicular epithelium that are committed to become squamous increase their compliance to deformation, which is also consistent with the measured softening of dorsal ectoderm cells [1]. In general, our results are therefore consistent with mechanical transitions that were measured with a force-exerting and lower frequency methodology, AFM.

Previous work using laser microdissection experiments showed that dorsal and mesodermal cells soften and neuroectodermal cells stiffen during gastrulation [8]: How can this be integrated with our result? Brillouin microscopy measures the stiffness of the entire cell volume with sub-cellular resolution; however, laser microdissection measures the cortical tensions (i.e., a force) along the apical surface of the epithelium, potentially averaging contributions from a larger field of cells. Furthermore, using Brillouin microscopy our measurements were made along the apical-basal axis of the cells, which is perpendicular to the axis probed by laser microdissection. Finally, we measured changes in mechanical properties at regular timepoints throughout gastrulation in the same specimen; on the contrary, laser microdissection can be performed once, during gastrulation, and therefore, it is not possible to assess the mechanical changes over time within the same specimen. These differences provide a possible explanation for the distinct mechanical properties of dorso-ventral embryonic cell populations between both studies and highlight the usefulness of complementary studies to comprehend the complex mechanics of three-dimensional shape acquisition fully.

In pre-cellularising *Drosophila* embryos, the elastic modulus of the cytoplasm is determined by F-actin [14]. But the increase in Brillouin shift in the furrow-forming cells cannot be fully explained by the actin cytoskeleton, because it does not colocalize fully with the known subcellular distribution of F-actin and actin-binding proteins in central mesodermal cells. We show that MTs predominantly contribute to cell mechanical properties in the central mesoderm, which is consistent with described roles of MTs in other instances of cell shape homeostasis [28]. MTs have properties that support their function as mechano-effectors: high persistent length [48], dynamic instability [49] and differential stiffness [50, 51]. In particular, dynamic MTs are stiffer than stable MTs, which enhance their flexibility through acetylation [50]. Previous work has shown that MTs in the lateral-basal compartment of mesodermal cells are acetylated [28], whereas sub-apical MTs are not acetylated and remain tyrosinated [28], suggesting that dynamic sub-apical MTs make the sub-apical compartment of central mesodermal cells stiff. The progressive alignment of sub-apical MTs with each other along the apical-basal axis is consistent with our measurements of an increasing Brillouin shift. One possible interpretation is that sub-apical MTs counteract the compression along the apical-basal axis to help the progression of folding.

Although we note that the physical toy model of VFF we developed does not simulate the Brillouin scattering process itself and therefore model stiffness values (a.u.) and frequency shifts (GHz) can not be directly compared, it corroborates our Brillouin measurements and highlights those localised changes in cellular material properties –which are often assumed to be constant during a morphological process-can facilitate the progress of folding events. The model predicts that only changes in sub-apical regions of the epithelial cells affect furrow formation (Fig. 5d). This fits well with our observations that Brillouin shifts and MT reorganization are restricted to that compartment, as well as with previous measurements in cellularising *Drosophila* embryos that showed a heterogeneous Young’s modulus along the pre-blastoderm cytoplasm [14]. While experimental evidence and previous models ascribe a driving function to apical constriction in furrow formation, our model showed that this process is contingent on the longitudinal stiffness of central cells (Fig. 5e’). A stiffer mesoderm along the apical-basal axis correlates with deeper furrows, but the model predicts better outcomes if cells are initially softer and stiffen over time, again a phenomenon we also see in the Brillouin measurements. This observation of spatially restricted cell stiffening therefore fits with a model in which cell-shape-driven morphogenetic events require cells to behave not as a uniform and constant mechanical unit.

We therefore propose a model in which the observed changes of mechanical properties of the mesoderm result from the rearrangement of microtubules, mediated by the apical constriction and elongation of central mesodermal cells. The organisation of microtubules is in the first instance determined by the geometry of the compartment in which they grow, with microtubules becoming more organised when the compartment elongates [35]. Thus, the space generated between the nuclei and the apical side of the cells during the apical constriction of central mesodermal cells could enable an initial increase in sub-apical MT length and alignment of MTs both with the apical-basal axis and with each other. Because in Brillouin microscopy, stiffness derives from the longitudinal modulus, measurements were mostly conducted along the cellular apical-basal axis, and thus, the increasingly aligned MTs within the sub-apical compartment Fig. 4c-e) are good candidates to dynamically stiffen mesodermal cells, which in turn would facilitate fold formation.

We describe different mechanical dynamics for the mesoderm and the ectoderm (Figs 1,2.). Although both ectodermal cell subpopulations, the neuroectoderm and the dorsal ectoderm differ in biological qualities that could affect cell mechanics, such as the accumulation of actomyosin and the localisation of adherens junctions [5, 28] and Cheerio/Filamin (Fig. 3c, top), we found similar mechanical dynamics among cells in the neuroectoderm and the dorsal ectoderm. Therefore, from a mechanical standpoint, we have identified two distinct domains along the DV axis: the mesoderm and the whole ectoderm. A genetic mechanism that could assign different mechanical properties to mesoderm and the whole ectoderm must rely on the mesodermal determinant Snail [52]. However, it is unclear what are the possible mechanoeffectors that may be regulated by Snail in the mesoderm. A recent proteomic and phosphoproteomic study of DV cell populations at gastrulation stage has found networks of proteins and phosphoproteins whose abundance is increased either in the whole ectoderm or in the mesoderm [28]. Future work will investigate the role of these regulated proteins and phosphoproteins in determining the mechanical dynamics of the ectoderm.

Altogether, our results show that cells can adjust their material properties within minutes to optimally drive morphogenesis, and highlights the potential of Brillouin microscopy as a powerful tool for investigating the mechanical aspects of cell shape behaviors and their relationship to tissue dynamics in vivo. Future work will determine the exact role of microtubules and their binding proteins in determining the evolving mechanical properties during tissue folding.

## Acknowledgements

We thank N.H. Brown, N. Bulgakova, S. De Renzis, S. Huelsmann, K. Roeper and Y-1. C. Wang for reagents and fly stocks. N. Lawrence and the Gurdon Institute Imaging Facility for help with 3D-SIM imaging, EMBL Advanced Light Microscopy Facility (ALMF), in particular 2. M. Lampe for assistance with Spinning Disk microscopy and image deconvolution, and CECAD Cologne Imaging Facility for continuous support. We acknowledge the computing resources provided by North Carolina State University High Performance Computing Services Core Facility (RRID:SCR_022168). Flybase was used throughout the project and is gratefully acknowledged. We thank E. Vogelsang for assistance with physical cross-sections of fixed embryos. We also thank C.J. Chan, N. Petridou, K. Prummel and Y.-C. Wang, for critical discussion and reading of the manuscript. Flybase was used throughout the project and is gratefully acknowledged. R.P. acknowledges support of an ERC Consolidator Grant (no. 864027, Brillouin4Life), and the German Center for Lung Research (DZL). This work was also supported by funding from the European Molecular Biology Organisation (EMBO), the University of Cologne and the Deutsche Forschungsgemeinschaft (grant DFG LE 546/12). This work was supported by the European Molecular Biology Laboratory (EMBL) and the North Carolina State University.

## Author contributions

J.M.G. conceived and designed the study as well as the analyses of processed Brillouin data and wrote the manuscript. C.B. and J.M.G collected the data and C.B. performed the analyses of raw Brillouin data. A.T. and J.M.B. developed and implemented the *in silico* model. M.L. contributed to the biological interpretation and discussion of results and aided in the selection of mutants. R.P. contributed to study and data analysis design, and its physical interpretation, acquired funding and supervised the project. All authors provided critical feedback on all aspects of the research, and contributed to the final manuscript.

## Materials and Methods

### *Drosophila* genetics, embryo collections and live imaging mounting

#### Drosophila genetics

To visualise cell membranes and nuclei (Fig. 1b-h, Fig. 2a-c), we used transgenic lines expressing CaaX-eGFP [28] and His2B-mRFP (both stocks provided by Yu-Chiun Wang). To study the distribution of lipid droplets, we used a YFP protein-trap for the gene *lsd-2* (provided by Yu-Chiun Wang, source is *Drosophila* Kyoto Stock Center/DGGR-Kyoto: 115301) [53].

To analyse the colocalisation between actomyosin and the Brillouin shift during ventral furrow formation (Fig. 3a), we combined the sqh-sqh::mCherry transgene (Bloomington stock N°: 59024) to visualise non-muscle myosin with a utrophinABD-GFP transgene [54].

To study the distribution of actin binding proteins we used the following transgenic lines: GT{cher[mGFP6-1]}/TM6 [55, 56] a GFP knock-in that specifically labels the long isoform of Cheerio (provided by Sven Huelsmann), PBac{768.FSVS-0}Fim[CPTI003498] (DGGR-Kyoto: N°: 115478) a protein-trap for Fimbrin, PBac{754.P.FSVS-0}tsr[CPTI002237] (DGGR-Kyoto: N°: 115280) a protein-trap for Cofilin/Twinstar. For immunostainings of actin binding proteins, we used either *w^1118^* (Bloomington stock: 5905), or Gap43-mCherry [57] (provided by Stefano De Renzis), a transgenic line that labels cell membranes.

To study the microtubules and the centrosome position (Fig. 4a) we used a transgenic line (genomic construct) that labels the centriolar protein Asterless (pUbi-YFPasl^FL^) [33].

For microtubule perturbation experiments (Fig. 4f-g), EB1-GFP visualisation (Fig. 4c-e) and analyses of the Brillouin shift during dorsal fold formation (Fig. 3b), we used a transgenic line that combined Gap43-mCherry with EB1-GFP [34] (provided by Nick Brown) to label the plus ends of the microtubules.

To study the connection between the folding event and invagination with the measurement of an increase in the Brillouin shift we used the *twist^1^* [58] mutant allele *twi^1^*/Cyo,hb-GFPnls; 3X-CaaX-mScarlet [28]/TM6ref (provided by Yu-Chiun Wang).

#### Embryo collections

For all live imaging experiments (using either line-scan Brillouin, spinning disk or 2-photon microscopes), we synchronised egg collections for 1 hour. The collected embryos were allowed to develop a further hour at 25°C. The embryos on the agar plates were covered with halocarbon oil (27S, Sigma) to hand-select embryos at the cellularisation stage (stages 5a,b [23]) using a Zeiss binocular. Finally, embryos were dechorionated (standard bleach, 50% in water) for 90 seconds, washed in PBS 1X and mounted for microscopy.

For immunohistochemistry experiments, we synchronised egg collections for 2 hours, and the collected embryos were allowed to develop further for 2 hours, dechorionated and transferred to a glass vial for immediate fixation (see immunohistochemistry).

#### Live imaging

For Brillouin microscopy experiments, dechorionated embryos were oriented to target the corresponding cell population (dorsal, lateral or ventral) and glued (heptane glue) on the surface of a custom-built tool made of a plastic material: polyether ether ketone. The temperature of the imaging chamber was 21°C +/− 1.5°C.

For vertical mounting in combination with 2-photon microscopy, dechorionated embryos were glued (heptane glue) on their posterior end to 35mm glass-bottom petri dishes (MatTek, 10mm microwell, N° 1.5 coverglass), with their anterior side pointing up, and embryos were oriented vertically to acquire a dorso-ventral cross-sectional view [28]. Finally, the embryos were embedded in 1% low-point melting agarose (Sigma-Aldrich, A9414) in PBS added at ∼32°C, and the agarose was allowed to cool to RT (22°C).

For Spinning Disk microscopy, dechorionated embryos were oriented with their ventral side to be in contact with the glass bottom of the petri dish and 6 ml of PBS were added.

### Immunohistochemistry

#### Fixation procedures

To detect and visualise microtubules and actin binding proteins while preserving the overall cellular structures, a formaldehyde-methanol sequential fixation was performed. Dechorionated embryos were fixed in 10% formaldehyde (methanol free, 18814 Polysciences Inc.) in PBS:Heptane (1:1) for 20 min at room temperature (RT). To visualise microtubules, we used ice-cold methanol:heptane and embryos were stored for 24 hours at –20°C and rehydrated before use [35]. To visualise actin binding proteins, we devitellinised for 45 sec in 1:1 methanol:heptane (RT methanol).

#### Antibody staining procedures

Rehydrated embryos were blocked for 2 h in 2% BSA (B9000, NEB) in PBS with 0.3% Triton X-100 (T9284, Sigma). Primary antibody incubations were done overnight at 4°C. Primary antibodies used were: mouse anti α-tubulin 1:1000 (T6199, clone 6-11B-1, Sigma), rabbit anti Snail [16] 1:500, rabbit anti GFP 1:1000 (ab290, Abcam), mouse anti FITC-GFP 1:250 (ab6662, Abcam), anti E-cadherin 1:50 (clone DCAD2, DSHB), anti Profilin/Chic (clone Chi-1J 1:50, DSHB), anti discs-large 1:50 (clone 4F3, DSHB) and anti α-Actinin 1:100 (ab50599, Abcam). Incubations with secondary antibodies were performed for 2 hours at RT. Alexa Fluor 488, 568, 594 and 647-coupled secondary antibodies were used at 1:600 (488: goat anti mouse IgG ab150117, Abcam– and goat anti rabbit IgG ab150081, Abcam; 568: goat anti rat IgG JacksonImmunoResearch, 115-295-166-; 594: goat anti mouse IgG ab150120, Abcam; 647: Goat anti rat IgG 112-605-167, JacksonImmunoResearch and goat anti mouse IgG JacksonImmunoResearch, 115-295-166).

#### Preparation of physical cross-sections

Physical cross-sections of embryos were made as previously described [28]. Embryos that had been immunostained were embedded in Fluoromount G (SouthernBiotech 0100-01) and visually inspected to identify embryos at stage 6 (gastrulation [23]). The selected embryos were sectioned manually with a 27G injection needle at approximately 50% embryo length and slices were mounted on the sectioned side.

### Confocal, spinning disk and 3D-SIM super-resolution image acquisition

Live imaging of CaaX-eGFP;H2B-mRFP embryos (Supplementary Figure 1a and Supplementary Video 1) was performed with a Zeiss LSM780 NLO 2-photon microscope with a Plan-Apochromat 63x objective (NA 1.4, oil, DIC M27) at a room temperature of 22°C. We acquired 4 slices (z-slice size = 1,1um) at approximately 180 μm from the posterior end of the embryo every 2 minutes. Supplementary Video 1 was generated by max-projecting 3 consecutive slices.

Images in Figures 3c and 4a were acquired with a Leica SP8 microscope equipped with a supercontinuum laser. Gated detection on HyD detectors was used for each channel using a Plan-Apochromat 63x oil (NA 1.4) objective at 22°C, with a z-slice size of 0.3 μm. Acquired volumes used in Figure 3c were max-projected (along the z-axis) for a range of approximately 1.5 μm (5 slices); volumes used in Figure 4a were max-projected (along z-axis) for a range of approximately 3 μm (10 slices).

Live imaging of EB1-GFP embryos (Fig. 4c-e and Supplementary Video 11) was performed with an Olympus IXplore SpinSR, with a pinhole size of 50 μm, using the C488-561 dual filter and equipped with 2x ORCA-FLASH 4.0v3 cameras. For acquisition we used a 100X UPLSAPO 100 XS objective (NA 1,35) with silicon immersion. Acquired volumes were YZ resliced and max-projected over 20 consecutive slices. Imaging was performed at a room temperature of 22°C. The acquired raw volumes for each embryo were deconvolved using Huygens Professional 23.1.

Images in Figure 4b were acquired using a super resolution Deltavision OMX 3D-SIM (3D-SIM) V3 BLAZE from Applied Precision/GE Healthcare. Deltavision OMX 3D-SIM System V3 BLAZE is equipped with 3 sCMOS cameras, 405, 488 and 592.5 nm diode laser illumination, an Olympus Plan Apo 60X 1.42 numerical aperture (NA) oil objective, and standard excitation and emission filter sets. Imaging of each channel was done sequentially using three angles and five phase shifts of the illumination pattern. The refractive index of the immersion oil (Cargille) was 1.516. Acquired volumes were max-projected (along the z axis) for a range of 9 μm (30 slices). Imaging was performed at a room temperature of 20°C.

### Brillouin microscopy

The microscope combines a self-built fluorescence light-sheet and a confocal line Brillouin modality as described in detail in Bevilacqua et al [16]. In brief, two water immersion objectives (Nikon MRD07420, 40x 0.8NA) are mounted at 90 degrees. For the fluorescence modality, one of the two objectives generates a static light sheet of approx. 200 μm width and 4 μm thickness and the fluorescence light is detected by the other objective and imaged on a sCMOS (Zyla, Andor). For the Brillouin modality a confocal line is generated by a cylindrical lens in combination with the objective, while the Brillouin back-scattered light is collected by the same objective and sent to a custom-built Brillouin spectrometer. Note that the lenses CL_2_ and CL_3_ in Bevilacqua et al. [16], are substituted by a single spherical lens of 150 mm, giving a pixel size along the line of 1.1μm instead of 0.7μm. The total laser power on the sample was kept below ∼22mW. The embryos were glued on a 3D-printed tool with a cut at 45 degrees, to facilitate the orientation of the embryos with respect to the detection objective. The tool was connected to a 3-axes translational stage and immersed directly in PBS. For the permeabilized embryos a 12.7 μm thick FEP foil was placed between the embryos and the detection objective to reduce the volume of PBS in contact with the embryos (see below: Microtubule perturbation using Colcemid).

### Microtubule perturbation using Colcemid

Synchronised egg collections of 45’ were allowed to develop for 2 hours at 25°C. Embryos were dechorionated in 50% bleach in water. We removed the the wax layer that covers the vitelline membrane [59] by fully immersing embryos in a permeabilising solution, i.e. EPS [59], 1:10 in PBS for 3 minutes. The EPS solution consists of 90% D-Lemonene (Sigma-Aldrich, 183164), 5% cocamide DEA (Ninol 11CM, Stepan Chemical, Northfield, Illinois) and 5% ethoxylated alcohol (Bio-Soft 1-7, Stepan Chemical, Northfield, Illinois). Next, embryos were promptly washed first in Triton 0.3% in PBS and subsequently in PBS only. Permeabilised embryos were visually inspected using a Zeiss stereoscope, and based on morphological criteria [23] embryos in stage 5a [23] were selected and mounted. Finally, mounted embryos were placed within the Brillouin microscope incubation chamber filled with either PBS or a solution of 1mg/ml Colcemid in PBS (Demecolcine, Sigma-Aldrich, D7385). Embryos that were in mid to late cellularization (membrane growth was visualised using the membrane marker Gap43m-Cherry) at the moment of initiation of the incubation with Colcemid were imaged throughout the gastrulation stage. Five independent experiments, each experiment with the acquisition of one embryo, were performed each for control (permeabilized + PBS) and treated embryos (permeabilised + 1mg/ml Colcemid in PBS). Imaging quantitation and statistical analyses were performed as described below.

### Image quantification

Image analyses were performed using ImageJ/Fiji version 2.14.0/1.54f and specifically for Fig. 4d,e, we also used Matlab R2019b (see below: Quantification of the alignment of microtubules with each other).

Masks were hand-generated in ImageJ/Fiji to filter the Brillouin signal of target tissue areas. For all quantitations with the exception of EB1-GFP analyses (Fig. 4c-e, see below: Quantification of the alignment of microtubules with each other), masks were generated based on the signal of a membrane marker (CaaX-eGFP or Gap43-mCherry) using the ‘polygon selections’ tool. We extracted shape parameters of the mask itself and values of the Brillouin shift signal within the region of interest (ROI) that was enclosed by the mask with the ImageJ/Fiji ‘measure’ tool. The shape parameters of the masks that were estimated were the major and short axis lengths and the circularity values. From the filtered Brillouin shift values, we estimated the median, mean, standard deviation, maximum and minimum Brillouin shift values. Mask generation and the above quantifications were performed on single slices from YZ resliced raw volumes.

#### Mask generation and measurements within region of interest

For the results shown in Fig. 1b,c, a mask to segment 16 cells with its centre in the ventral midline was generated manually for each timepoint from the onset of gastrulation to mesoderm invagination using the CaaX-eGFP signal for the identification of cell boundaries. Once the mesoderm was invaginated, the CaaX-eGFP signal corresponding to cells deep inside the embryo scattered, difficulting the delimitation between the interior-most cells and the yolk for the generation of masks. To circumvent this problem, we generated a mask by thresholding the CaaX-eGFP signal at the time-point the folding is initiated. The threshold was set to filter out any signal that does not correspond to membranes in any timepoint subsequent to the invagination of the mesoderm. Once the mask was generated, it was overlaid onto the CaaX-eGFP signal and used as a guide to delineate a boundary between the invaginated mesoderm and the yolk for the generation of masks corresponding to the mesoderm cells undergoing EMT.

For the results shown in Fig. 1d,e, a mask consisting of 6 cells positioned at a distance of 11-13 cells dorsally to the boundary of the ventral fold (counted using the nuclear signal from the H2Bav-mRFP marker) was generated for each time-point from the onset of gastrulation to the point where the ventral displacement of neuroectoderm cells is complete.

For the results shown in Fig. 1f,g we generated masks for 6 cells that are positioned in the dorsal ectoderm for each timepoint from the end of cellularisation (when the membrane signal stops growing towards the yolk, Supplementary Video 2) until squamous morphogenesis generates stretched cells. We excluded from our analyses the presumptive amnioserosa, which we considered to be 6 cells wide around the dorsal-midline (Fig. 2c).

For the results shown in Fig. 2a, a mask was generated for each cell of the mesoderm (along a 16-cell field with its centre in the ventral midline), from the onset of gastrulation until the initiation of the folding event.

For the results shown in Fig. 2c-g, a mask was generated for a single cell for each time-point from the end of cellularisation until the squamous shape had been acquired. We wanted to analyse the correlation between the Brillouin-shift and the length of the apical-basal axis during squamous morphogenesis, using the major and minor axes measurement derived from fitting ellipses with ImageJ/Fiji together with the circularity measurement (Supplementary Fig. 2a; b: 1-3). When the cells flatten during squamous morphogenesis, the apical-basal axis, which was initially the major axis of the fitted ellipse becomes the minor axis when the cell goes through an isotropic shape (Supplementary Fig. 2b: 3). The perpendicular axis to the apical-basal axis, termed ‘stretching axis’, which was initially the minor axis of the fitted ellipse, becomes the major axis after the cell goes through an isotropic shape (Supplementary Fig. 2b: 3). The time point when cell masks become isotropic is the time point with the largest circularity value, this is, when cells are approximately cuboidal (Supplementary Fig. 2b: 4-5). Next, we measured in each cell the apical-basal axis by combining the major cell axis measurements before the time point the cell acquires a max circularity value, with the minor axis measurements after that time point; the stretching axis was assembled combining the minor axis before the max circularity timepoint, with the major axis measurements after that timepoint (Supplementary Fig. 2b: 6). Finally, for each cell we generated time course tables for the cross-correlation of the median Brillouin shift measured within each cell with the corresponding shape parameters (apical-basal and stretching axes length in μm).

To plot an average time course of the cell-by-cell quantification of the Brillouin shift and the length of the apical-basal and stretching axes (Fig. 2f), we filtered those cells that acquired the largest circularity value between 00:42:20 and 01:07:44 hours, corresponding to the 10th and 90th percentile of the distribution of maximum circularity times, leading to the inclusion of 7/9 cells. The maximum circularity value was measured at a median of 00:53:22 min, and this value was used in Fig. 2f to set the boundary between phase 1 and phase 2.

For the results shown in Fig. 4c,d we median-projected 3 consecutive slices of the XZ resliced volume, and generated masks for 6 cells for central mesodermal cells (with their centre on the ventral midline), for each time point from the onset of gastrulation until the mesoderm had invaginated (permeabilised embryos in PBS/control) or mesoderm cells had undergone EMT on the surface of the embryo (permeabilised embryos in 1 mg/ml Colcemid in PBS).

### Subcellular localisation of high-Brillouin shift

To filter the largest 4% Brillouin shift values, we applied a ‘default’ threshold using ImageJ/Fiji on the Brillouin shift maps, using the end of cellularisation as a reference timepoint to apply the threshold.

### Quantification of the alignment of microtubules with each other

For each EB1-GFP-expressing embryo (N = 3) acquired using spinning disk microscopy, we YZ re-sliced the deconvolved raw volume, and generated 5 max-projections of 20 consecutive slices each. Next, we generated rectangular masks (covering a tissue section of 73.3 μm^2^) that were placed within the sub-apical compartment of central mesodermal cells in each time point of each max-projected movie of each embryo. Thus, for all analysed embryos, across all timepoints, we utilised a mask of the same geometry and the same size. The rectangular masks were used to filter the EB1-GFP signal within the sub-apical compartment of central mesodermal cells.

We analysed the alignment of the filtered EB1-GFP signal with a published Matlab (R2019b) script that analyses the mean direction of the microtubule signal (either α-Tubulin immunostainings or EB1-GFP live imaging) and the dispersion of the signal around the estimated mean microtubule direction, which was termed ‘MTSD’ [28]. Using this Matlab script, we obtained for each embryo 5 MTSD values (one for each max-projected movie) for every analysed time point. The 5 MTSD values obtained for each timepoint were averaged, resulting in a mean MTSD and standard error of the mean per timepoint and per embryo (Fig. 4d).

### Statistical analyses

All statistical analyses were performed in Prism Graphpad 10.2.3. Before each analysis, normality testing was performed using both Shapiro-Wilk test and Normal QQ plots, and based on both tests the decision to conduct parametric or non-parametric tests was made. In Fig. 1c,e,f we conducted a one-way ANOVA (parametric, matched measures); in Fig. 1h we conducted a simple linear regression for the pooled neuroectoderm and the ectoderm between timepoints –0:04:16 hour (end of cellularisation) and 01:01:23 hour (ongoing squamous morphogenesis) and tested whether the resulting slope and intersects were significantly different when separate linear regressions (i.e. for the neuroectoderm and ectoderm separately) were conducted; in Fig. 2g we conducted a Pearson correlation between the apical-basal or stretching axes length and the median Brilloiun shift for each quantified cell from the end of cellularisation (t=0’) until the last quantified time point (squamous morphogenesis ongoing); in Fig. 4e a non-parametric ANOVA Friedman test was conducted; in Fig. 4g a non-parametric Mann-Whitney test was conducted. We applied False Discovery Rate (two-stage step-up method of Benjamini, Krieger and Yekutieli; q < 0.05) to correct for multiple comparisons. For all statistical tests significance was set at 5% (p < 0.05).

### Physical model of gastrulation

The physical model of *Drosophila* VFF was developed within the Cellular Potts (Glazier-Graner-Hogeweg) modelling framework [37], where individual cells and their internal compartments are represented by a collection of lattice sites on a fixed grid which evolves in time according to a Metropolis algorithm that minimises an effective energy that describes cell properties and behaviours (see SI for a detailed description of the model). The model was developed using the CompuCell3D (CC3D) simulation software [60], and the simulations ran on a computer cluster using a set of custom developed python scripts to efficiently explore the parameter space. The code for the simulations is publicly available at https://github.com/abhisha-ramesh/Gastrulation_Model.

