## Supplementary Information for "Highly dynamic mechanical transitions in embryonic cell populations during *Drosophila* gastrulation"

Supplementary Figure 1

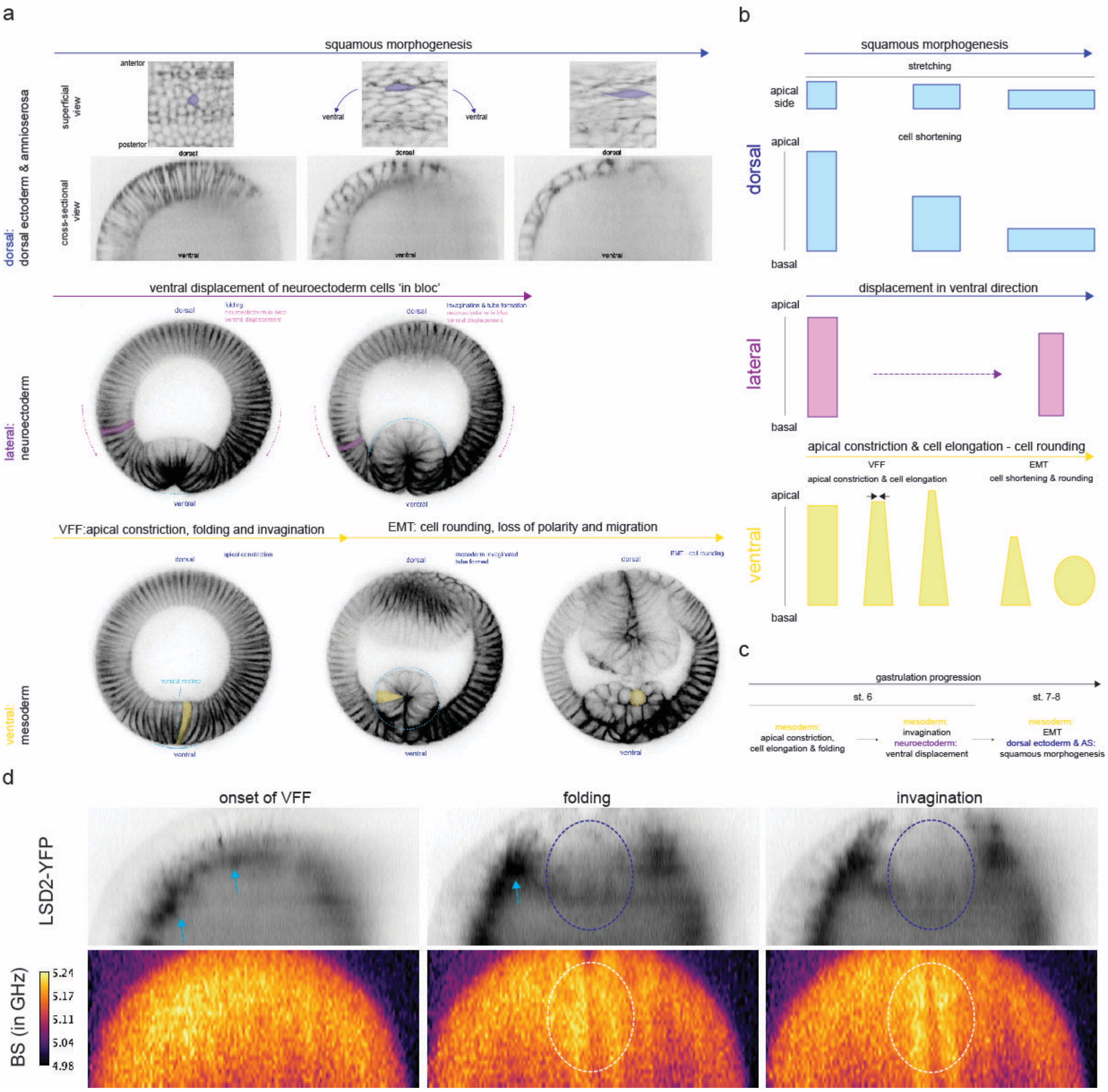

**Cell shape behaviours in cells along the dorso-ventral axis during *Drosophila* gastrulation.**

**a.** Description of cell shape behaviours within each DV cell population. Top: dorsal side of the embryo, showing squamous morphogenesis in dorsal ectoderm and amnioserosa (AS) cells. The process of squamous morphogenesis leads to the stretching of cells (along the embryonic DV axis, anterior is top, blue shaded cell) and the shortening of cells (along the cellular apical-basal axis). Lateral and Ventral: 2-photon cross-sectional imaging of a *Drosophila* embryo showing cell shape behaviours within the Neuroectoderm (lateral, magenta shaded cell) and Mesoderm (ventral, yellow shaded cells). Neuroectoderm cells display little changes in their cellular geometry; Mesoderm cells undergo apical constriction and cell lengthening (during VFF, left panel), which is followed by cell shortening and rounding during EMT (centre and right panels respectively). In all cross-sectional panels, dorsal is top, ventral is bottom.

**b.** Illustration that schematises the changes in cellular geometry in the dorsal, lateral and ventral side of the embryo.

**c.** Coordination of morphogenetic events within DV cell populations (yellow: mesoderm; magenta: neuroectoderm; blue: dorsal ectoderm and AS) during gastrulation (starting in stage 6, and continued during stage 7 and 8), based on work by Rauzi et al. [1].

**d.** Colocalisation between a protein trap for *lsd-2*, lsd2-YFP (top panel), and the transient increase in Brillouin shift (BS in GHz, bottom panel) that is measured within the mesoderm during VFF. Lsd2 associates with lipid droplets and vesicles. Area of the mesoderm encircled with a blue and white-dashed line shows increase in the Brillouin shift during folding and invagination. Images are average projections of two consecutive YZ re-slices of the original volume. Note that Lsd2-YFP puncta do not colocalise with the high Brillouin shift in central mesoderm.

### Supplementary Figure 2

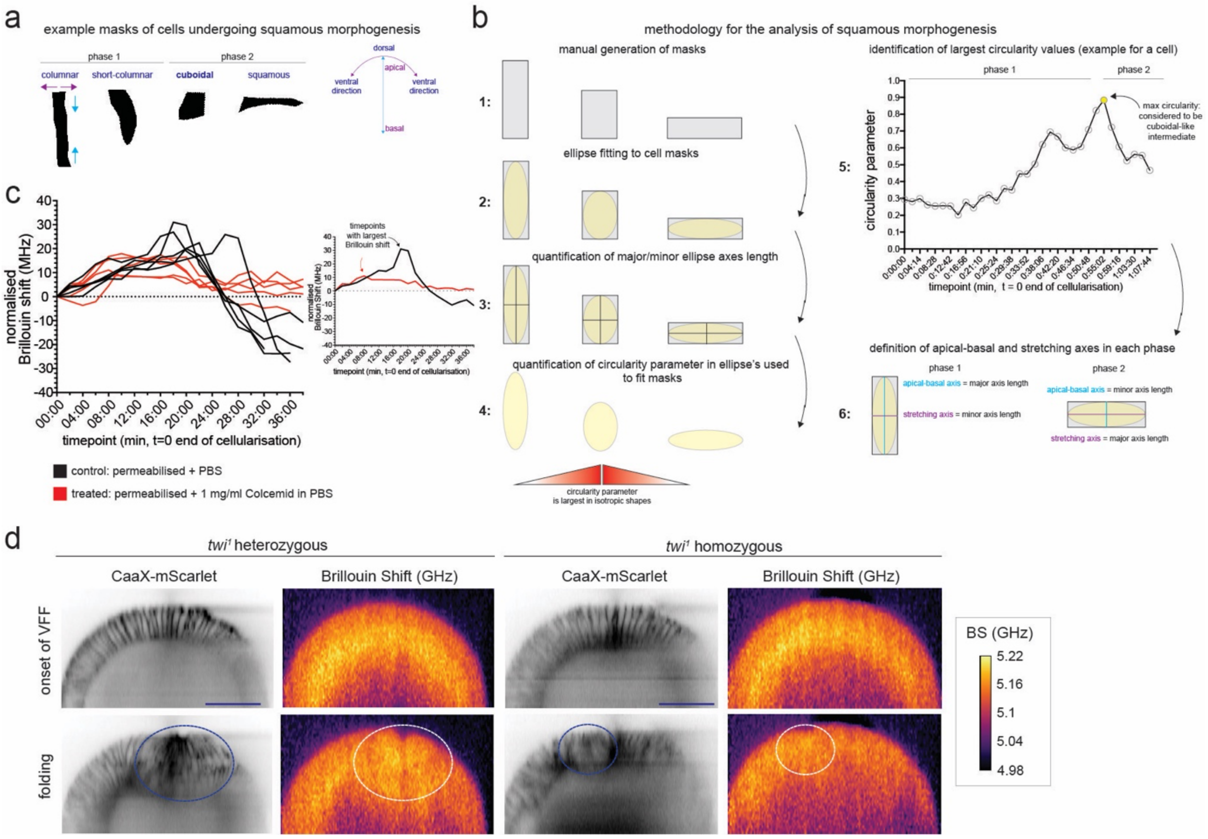

**Cell shape behaviours in cells along the dorso-ventral axis during *Drosophila* gastrulation.**

**a.** Description of progressive changes in cell geometry during the squamous morphogenesis of dorsal cells (dorsal ectoderm and AS) at gastrulation stage. Shapes are examples of manually-generated cell masks of a dorsal cell of an embryo at gastrulation stage. Right panel shows the axes in which changes in cell geometry occur: cell apical-basal axis (cyan) and along the perimeter of the embryo in the dorso-ventral direction (magenta). During squamous morphogenesis cells shorten along the apical-basal axis and stretch in the ventral direction. We defined two phases during squamous morphogenesis, *phase 1* prior to the formation of the 'cuboidal' intermediate, and *phase 2*, subsequently.

**b.** Illustration that describes key steps (1-6) of the methodology used to analyse the cellular shapes during squamous morphogenesis. Cells that could be followed in a single reslice of the acquired volume from the end of cellularisation until cells became squamous were manually segmented (step 1). The measurements of the major and minor axes length (step 2-3) were used to quantify the apical-basal (cyan) and stretching axes (magenta, perpendicular to apical-basal axis). To detect the 'cuboidal' intermediate we used the circularity parameter, which is larger when cells are increasingly isotropic (steps 4-5, yellow dot in circularity parameter timelapse). The cell mask with the maximum circularity value was considered to represent the 'cuboidal' intermediate, and the boundary between phase 1 and 2. During phase 1, the major axis length was used to measure the apical-basal axis and the minor axis was used to measure the stretching axis. During phase 2, the major axis length was used to measure the stretching axis, and the minor axis was used to measure the apical-basal axis.
**c.** Quantification of the BS within 6 central mesoderm cells (3 cells on each side of the ventral midline, see Supplementary Fig. 1a, bottom panel and Supplementary Movie 1) during VFF in permeabilised embryos treated with Colcemid 1 mg/ml in PBS or left untreated (control, PBS). Each line is a single embryo and an independent experiment (N=5 for each condition). Control embryos are shown with black lines, Colcemid-treated embryos are shown with red lines. Quantifications were performed in median projections of three consecutive YZ reslices of the acquired raw volume. BS was normalised to the onset of VFF (t = 0'). Right inset shows a control and a Colcemid-treated embryo, and black/grey arrows indicate the maximum BS value that was used to perform the quantification of the effect of Colcemid treatment on the transient increase in BS during VFF (Fig. 4f,g).
**d.** Brillouin shift maps in *twist* (*$twi^1$,* right panel) mutant embryos during VFF. *twist* mutants engage a smaller number of cells that in control embryos (*$twi^1$*/+, left panel), form a smaller fold and fail to invaginate the mesoderm; note the transient high Brillouin shift is strongly reduced in *$twi^1$*. Blue and white dashed-line circles indicate the area of the blastoderm engaged in the folding event. Scale bar is 50 μm.

### Supplementary Figure 3

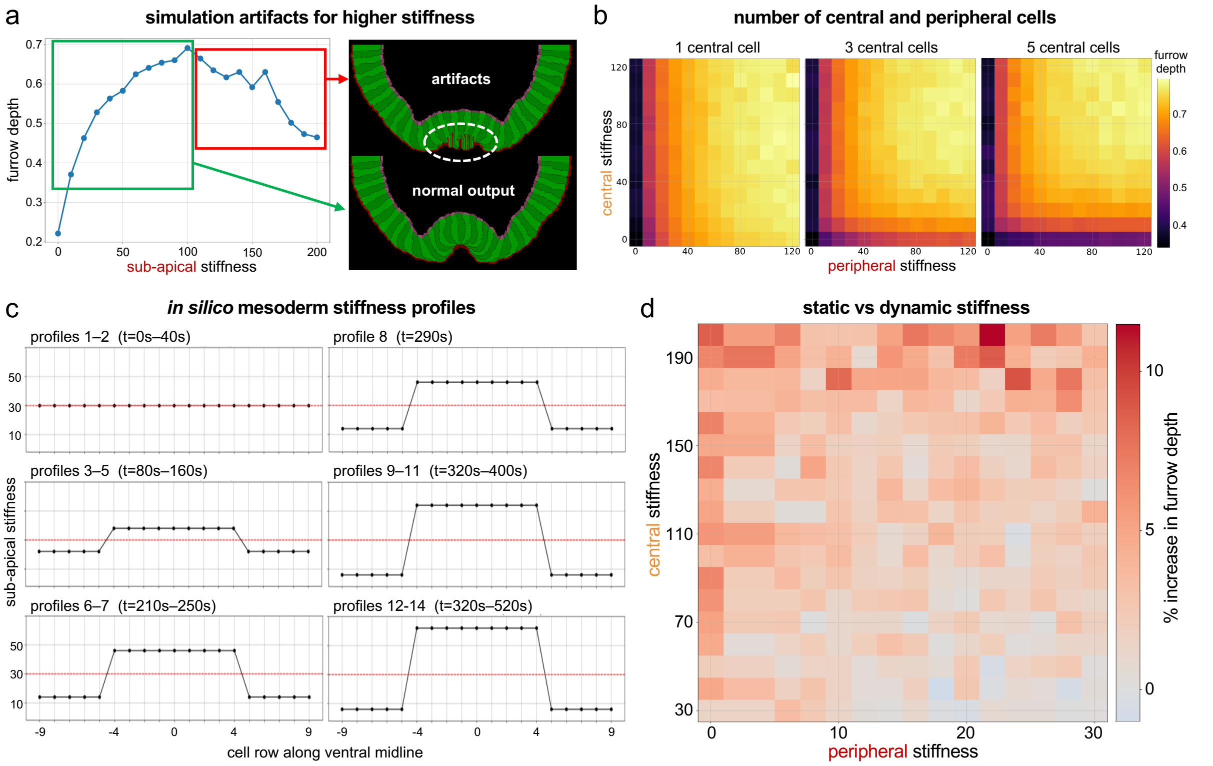

**Complementary and supporting results for the physical model of VFF.**

**a.** Furrow depth as a function of sub-apical longitudinal stiffness (and sub-basal stiffness set to 50 a.u.). For low values of longitudinal stiffness (green box) the model behaves normal and furrow depth correlates with higher stiffness. When stiffness is increased beyond 100 a.u. (red box) the model starts presenting artefacts and results can no longer be trusted.

**b.** Effect of different definitions of central vs peripheral cells in the results about the role of central vs peripheral cell stiffness in furrow formation. Results remain the same when a similar definition for the number of central cells as in Fig. 5e is used.

**c.** Set of 9 sub-apical longitudinal stiffness profiles along the ventral midline used in our simulations (Fig. 5f). Cells start with uniform stiffness throughout the mesoderm, with an initial softening of the peripheral cells (profiles 3 to 4), followed by a stiffening of the central cells (profiles 5 to 9). The maximum and minimum values of central and peripheral cell stiffness is different for each simulation (Fig. 5e).

**d.** Quantitative comparison between static (Fig. 5e) and dynamic stiffness (Fig. 5f) on furrow formation. Red colours represent an increased furrow depth for dynamic stiffness compared to the static case.

### Supplementary Video 1

Cross-sectional live imaging of a *Drosophila* embryo from the onset of gastrulation (stage 5b) until the initiation of mesodermal EMT (stage 8). Key morphogenetic events that occur in cells distributed along the DV axis are indicated in the upper right corner. Membranes were labelled with a CaaX-eGFP (grayscale) transgene. The depth of the furrow increases with the progression of VFF. Imaging was conducted in a 2-photon microscope, and embryos were mounted in agar, vertically with respect to the glass bottom of the petri dish. The imaging was conducted approximately 180 μm from the posterior side of the embryo, and three consecutive planes were max-projected. Dorsal is top, ventral is bottom. Scale bars are 50 μm.

### Supplementary Video 2

Live imaging of the dorsal side of a *Drosophila* embryo describing the squamous morphogenesis of the dorsal ectoderm and the amnioserosa cell populations using our custom-built Selective Plane Illumination Microscopy (SPIM) microscope coupled to the LSBM. Membranes were labelled with CaaX-eGFP (magenta) and nuclei were labelled with His2B-mRFP (green) transgenes. Top left and bottom panels show a superficial (max-projection of 4 slices), in which the process of cellular stretching is appreciated. Top right panel shows the YZ resliced volume (1 slice), in which the changes in cellular morphology along the apical-basal axis can be seen. In top left and bottom panels anterior is left and ventral is top-bottom. In the top-right panel, dorsal is top. Scale bars are 50 μm.

### Supplementary Video 3

Brillouin shift (BS, bottom) maps of the ventral side of the embryo (ventral is top), showing the dynamic changes in mechanical properties of the mesoderm from the onset of VFF (stage 5b) through mesodermal EMT (early stage 8). Imaging was performed on living embryos carrying transgenes that label membranes (CaaX-eGFP, top) and nuclei (His2B-mRFP, centre). Video was assembled by YZ re-slicing the original raw volume and averaging two consecutive slices. Region shaded in yellow in CAAX-eGFP corresponds to the mesoderm cells that were used for quantification. Scale bar is 50 μm.

### Supplementary Video 4

Colocalisation between the transient, high Brillouin shift (BS, bottom) measured in the mesoderm during VFF and lipid droplets (top). Imaging was performed on a YFP protein-trap transgenic line for the gene *lsd-2*, whose product binds to lipid droplets. Video was assembled by YZ re-slicing the original volume and averaging two consecutive slices. The ventral side of the embryo is top. Note that Isd2-YFP puncta do not colocalise with the high Brillouin shift in central mesoderm. Scale bar is 50 μm.

### Supplementary Video 5

Brillouin shift (BS, bottom) maps of the lateral side of the embryo (lateral is top, ventral is left), showing the dynamic changes in mechanical properties of the neuroectoderm from the onset of VFF (stage 5b) until the neuroectoderm has fully displaced in the ventral (left) direction. Imaging was performed on living embryos carrying transgenes that label the membranes (CaaX-eGFP, top) and the nuclei (His2B-mRFP, centre). Video was assembled by YZ re-slicing the original volume and averaging two consecutive slices. Region shaded in yellow in CAAX-eGFP corresponds to the neuroectoderm cells that were used for quantification. Scale bar is 50 μm.

### Supplementary Video 6

Brillouin shift (BS, bottom) maps of the dorsal side of the embryo (dorsal is top), showing the dynamic changes in mechanical properties of the dorsal ectoderm and amnioserosa cells from the end of cellularisation (stage 5b) through the squamous morphogenesis of the dorsal ectoderm and amnioserosa (stage 7 and early stage 8). Imaging was performed on living embryos carrying transgenes that label the membranes (CaaX-eGFP, top) and the nuclei (His2B-mRFP, centre). Video was assembled by YZ re-slicing the original volume and averaging two consecutive slices. Region shaded in yellow in CAAX-eGFP corresponds to the ectoderm cells that were used for quantification. Scale bar is 50 μm.

### Supplementary Video 7

Brillouin shift (BS, bottom-left and bottom-right) maps of the ventral side of the embryo (ventral is top), showing the spatial distribution of pixels with the largest 4% Brillouin shift (thresholded signal in top-right panel; pixels enclosed by white line in bottom right panel) during VFF. Imaging was performed on living embryos carrying transgenes that label the membranes (CaaX-eGFP, magenta, top-left) and the nuclei (His2B-mRFP, cyan, top-right). Video was assembled by YZ re-slicing the original raw volume and averaging two consecutive slices. Region shaded in yellow in CAAX-eGFP corresponds to the ectoderm cells that were used for quantification. Dotted white line indicates the basal boundary of the growing sub-apical compartment. Scale bar is 50 μm.

### Supplementary Video 8

Colocalisation between the transient, high Brillouin shift (BS, bottom) measured in the central mesodermal cells during VFF, F-actin (labelled with a UtrophinABD-GFP transgene; grayscale; top) and non-muscle myosin light chain (labelled with a sqh-mCherry transgene; grayscale ; centre). Note the transient high Brillouin shift distributed across a larger tissue section than apical actomyosin. Video was assembled by YZ re-slicing the original volume and averaging two consecutive slices. Scale bar is 50 μm.

### Supplementary Video 9

Mid-sagittal Brillouin shift (BS, bottom) maps of the dorsal side of the embryo (dorsal is top, anterior is left), showing the dynamic changes in mechanical properties during dorsal fold formation (DFF) from the end of cellularisation (stage 5b) until the end of posterior midgut (PMG) invagination (stage 8). Imaging was performed on living embryos that carry a transgene that labels the membranes (Gap43-mCherry; grayscale; top). Video was assembled by YZ re-slicing the original volume and averaging two consecutive slices. Anterior and posterior folds position along the anterior-posterior axis are indicated by arrowheads during the progression of DFF. Scale bar is 50 μm.

### Supplementary Video 10

Brillouin shift (BS, bottom) maps of the ventral side of the embryo (ventral is top), showing the effect of Colcemid treatment (concentration: 1mg/ml in PBS; right panels) on the dynamic increase in BS measured within central mesodermal cells during VFF (PBS control, left panels; cells shaded in yellow). Imaging was performed on living embryos that carry a transgene that labels the membranes (Gap43-mCherry; grayscale; top). To deliver Colcemid, embryos were permeabilised (see Methods). Video was assembled by YZ re-slicing the original volume and averaging two consecutive slices. Region shaded in yellow in CAAX-eGFP corresponds to the mesoderm cells that were used for quantification. Scale bar is 50 μm.

### Supplementary Video 11

Live imaging of EB1-GFP (grayscale) in embryos undergoing VFF, using a spinning disk microscope. Ventral is top. The sub-apical compartment in central mesoderm is enclosed by yellow-dashed shapes, at the onset of VFF and during fold formation. Video was assembled by YZ re-slicing of the original volume and max-projecting 20 consecutive slices. Scale bar is 25 $\mu$m.

### Supplementary Video 12

Brillouin shift (BS, bottom) maps of the ventral side of the embryo (ventral is top), showing the effect of *twist* loss of function (*twi*$^{1}$ homozygous mutation; right panels) on the dynamic increase in BS measured within central mesodermal cells during VFF (*twi*$^{1}$ heterozygous mutation; left panels). Imaging was performed on living embryos that carry a transgene that labels the membranes (CaaX-mScarlet; grayscale; top). Video was assembled by YZ re-slicing the original volume and averaging 2 consecutive slices. Scale bar is 25 $\mu$m. Scale bar is 50 $\mu$m.

### Supplementary Video 13

Video of a simulation where the sub-apical regions of both central and peripheral cells are soft ($\lambda_{sA}$=0) leading to failure of ventral furrow formation. Sub-basal stiffness of all mesodermal cells is set to $\lambda_{sB}$=50. Video corresponds to the first panel of Fig. 5e'. Simulated time from 0 to 8.6min.

### Supplementary Video 14

Video of a simulation where the sub-apical regions of central cells are soft ($\lambda_{sA,c}$=0) leading to excess elongation along their apical-basal axis and failure of ventral furrow formation. Sub-apical stiffness of peripheral cells is set to $\lambda_{sA,p}$=90, and sub-basal stiffness of all mesodermal cells is set to $\lambda_{sB}$=50. Video corresponds to the second panel of Fig. 5e'. Simulated time from 0 to 8.6 min.

### Supplementary Video 15

Video of a simulation where the sub-apical region of central cells is stiff ($\lambda_{sA,c}$=110), but the peripheral cells are soft ($\lambda_{sA,p}$=0). The high deformability of the peripheral cells prevents the furrow from ingressing further (compare with Supplementary Video 16). Sub-basal stiffness of mesodermal cells is set to $\lambda_{sB}$=50. Video corresponds to the third panel of Fig. 5e'. Simulated time from 0 to 8.6min.

### Supplementary Video 16

Video of a simulation where the longitudinal stiffness of the sub-apical regions of both central and peripheral cells is high ($\lambda_{sA,c}$=110 and $\lambda_{sA,p}$=90) leading to a deeper ventral furrow. Sub-basal stiffness of mesodermal cells is set to $\lambda_{sB}$=50. Video corresponds to the last panel of Fig. 5e'. Simulated time from 0 to 8.6 min.

### Supplementary Note 1

Our physical model of *Drosophila* VFF was developed within the Cellular Potts (aka Glazier-Graner-Hogeweg) modelling framework [2] using the CompuCell3D (CC3D) simulation software [3]. In this modelling framework, biological cells and their subcellular compartments are spatially represented as a collection of lattice sites in a regular (cartesian) grid with the same ID. Our model consists of a 2D cross-section of the ventral half of the *Drosophila* embryo, consisting of 47 cells, including 19 mesodermal cells and 28 neuroectodermal cells (Fig. 5a). Each cell is composed of three compartments: **apical, core** and **basal**, reflecting the apical–basal polarity of blastoderm.

An effective energy equation ($H_{\mathrm{Total}}$) defines cell/domain properties and behaviours such as size, aspect-ratios, adhesion preferences and interactions with other cells (Eq. 1). Each of these properties is governed by an individual term in the overall effective energy equation: a volume constraint ($H_{\mathrm{V}}$) helps maintain the size and compressibility of cell compartments; a surface constraint ($H_{\mathrm{S}}$) helps maintain the perimeter of apical compartments; a contact energy ($H_{\mathrm{C}}$) specifies the relative strength of contact/adhesion between compartments from the same and/or different cells; and a spring-like intercellular force term ($H_{\mathrm{F}}$) linking compartments' centre-of-mass can be used to model apical constriction between mesodermal cells and sub-apical and sub-basal longitudinal stiffness of cells by linking apical-core and core-basal cell compartments, respectively.

(Eq. 1) $$H_{\mathrm{Total}} = H_{\mathrm{V}} + H_{\mathrm{S}} + H_{\mathrm{C}} + H_{\mathrm{F}}$$

The system evolves by a series of random lattice-site copy attempts where a lattice site "i" is randomly selected, and a neighbouring lattice site "j" is randomly selected within the 4th neighbour order of "i". If the two lattice sites belong to different cells or cell compartments, we evaluate the difference in energy ($\Delta H$) if the ID of site "i" is copied over site "j". If the effective energy is reduced ($\Delta H<0$) the copy is accepted; if it is increased then the copy attempt is accepted with a probability equal to:

(Eq. 2) $$\exp(-\Delta H/T)$$

where $T$ is a parameter used to specify the level of fluctuations in the system. A monte-carlo step (MCS) is defined as $N$ lattice-site copy attempts, where $N$ is the number of lattice sites in the grid. In our model 1000 MCS corresponds to 20 seconds.

The volume constraint is used to penalise small deviations of cell compartments volume (v) – which in a 2D simulation is defined as the number of lattice sites occupied by the cell compartment – from their target volume ($v_{\mathrm{t}}$):

(Eq. 3) $$H_{\mathrm{V}} = \lambda_{\mathrm{V}}(v-v_{\mathrm{t}})^2$$

where $\lambda_{\mathrm{V}}$ is parameter proportional to the bulk modulus of the cell compartment.

The surface constraint is used to penalise small deviations of cell compartments surface (s) – which in a 2D simulation is defined as the total perimeter that the occupied lattice sites of the cell compartment has – from their target surface ($s_{\mathrm{t}}$):

(Eq. 3) $$H_{\mathrm{S}} = \lambda_{\mathrm{S}}(s-s_{\mathrm{t}})^2$$

where $\lambda_S$ is a parameter setting the strength of the constraints. This constraint is only used for apical compartments to prevent their fragmentation during apical constriction.

The contact energy ($H_C$) is modelled as:

(Eq. 4) $$H_c = \sum_i \quad \sum_{j \neq i} \quad J(\sigma_i, \sigma_j)$$

where the second sum is over the 4th neighbourhood of site "i"; $\sigma_i$ and $\sigma_j$ are the type of cell compartments occupying sites "i" and "j", respectively; and J is a matrix containing the contact energy between compartments belonging to the same cell and different cells.

The spring-like intercellular force constraint ($H_F$) has the form of:

(Eq. 5) $$H_F = \lambda_F(d\text{-}d_t)^2$$

where $d$ is the distance between two cell compartments centre-of-mass (defined by the spatial distribution of the lattice sites it currently occupies in the grid); $d_t$ is the target distance between them; and $\lambda_F$ is the strength of the constraint. A different $H_F$, with its own parameters $\lambda_F$ and $d_t$ is defined for each pair of cell compartments that are linked by such a force.

Assuming a linear relation between the longitudinal stiffness and the longitudinal strain, we use (Eq. 5) to model the longitudinal stress/strain relation of the mesodermal cells along the apical-basal axis by linking their internal compartments:

(Eq. 6a) $$H_{sA} = \lambda_{sA}(d_{a\text{-}c}\text{-}d_{a\text{-}c,t})^2$$

(Eq. 6b) $$H_{sB} = \lambda_{sB}(d_{b\text{-}c}\text{-}d_{b\text{-}c,t})^2$$

where the $d_{a\text{-}c}$ and $d_{b\text{-}c}$ is the distance between the centre of mass of the apical and core compartments, and the basal and core compartments, respectively; $d_{a\text{-}c,t}$ and $d_{b\text{-}c,t}$ are the initial distance between those compartments at the start of the simulation; and $\lambda_{sA}$ and $\lambda_{sB}$ are parameters proportional to the longitudinal modulus of the sub-Apical and sub-Basal regions of the modelled cells (which we will now on simply refer as the "stiffness" of the compartments).

The contractile force generated as a result of medial-apical actomyosin accumulation is implemented in the model using spring-like forces that link the 19 central mesodermal cells at apical sides ($H_A$). This constraint is added to the effective energy as

(Eq. 7) $$H_A = (M/\langle S \rangle)(d\text{-}d_t)^2$$

where the $d$ is the distance between the centre of mass of the two apical compartments; $d_t$ is the target distance between these compartments (set to 2 lattice sites); and the ratio M/<S> is the strength of the contractile force, where M is the time-varying myosin levels and <S> is the average apical surface area with the medium of the two linked cells. We utilised measured myosin levels dynamics within the mesoderm [4] to construct the time-varying profiles of the myosin levels in our simulations and to set the time stamp of the simulations.

Furrow depth in our simulations is measured as the normalised distance between the apical-most coordinate of the central mesodermal cell and the lowest point of the mesoderm from the initial condition in units of initial cell height in the apical-basal axis (Fig. 5a). All measurements of furrow depth reported in the parameters maps of Figs. 5d-e are averages of 10 to 30 simulations. For most of our analysis longitudinal cell stiffness was varied over a range from 0 to about 110 (values in arbitrary units), with higher values always leading to deeper furrows. In our model, increasing stiffness beyond those values lead to simulation artefacts such as shown in Supplementary Fig. 3a and we chose to exclude those cases from our analysis. Because of that we were unable to comment on how higher levels of stiffness may also prevent furrow formation by not allowing for any cell deformation.

For the analysis of the impact of stiffness gradient along the ventral mesoderm we split our modelled mesoderm between 9 central cells and 10 peripheral cells (5 on each side). This was done to simplify our analysis and the choice of 9 central cells was done to reflect the measured point of inflection in stiffness value measured in the experiments. Choosing a slight lower number of central cells (5) had very little quantitative effect in our results (Supplementary Fig. 3b last panel). It is only when the number of central cells is defined to be lower than 3 that we see a drastic qualitative difference in our results (Supplementary Fig. 3b first panel).

For the simulations with varying cell stiffness over time, we assumed constant sub-basal stiffness ($\lambda_{sB}$=50) for all cells and the same initial value of sub-apical stiffness for central and peripheral cells ($\lambda_{sA}$=30). Choosing a higher(lower) starting value for the cell stiffness leads to less(more) prominent results regarding the gain of furrow depth for dynamic changes in stiffness and also for the static vs dynamic comparison.

For the quantitative comparison between static and dynamic stiffness (Supp Fig 3d) we calculated the % increase in furrow depth from the static case ($F_S$, when the cells already start with the central and peripheral stiffness indicated by the coordinate values) and the dynamic case ($F_D$, when all cells start with a uniform stiffness of 30, and modify their central and peripheral cell stiffness over time until the value indicated by the grid coordinates). The formula used was 100%x($F_D$-$F_S$)/$F_S$.

Table S1: List of parameters used in the model

| Parameter | Name | Value |
|---|---|---|
| T | **CPM fluctuation amplitude** | 25 |
| n_copy | **Neighbour range for lattice site copy attempts** | 2 |
| n_contact | **Neighbour range for contact energy calculations** | 4 |
| t | **Total time** | 26000 MCS |
| *λ_v* | **Strength of Volume Constraint** | |
| | Apical(mesoderm) | 2 |
| | Apical(neuroectoderm) | 100 |
| | Core(mesoderm) | 10 |
| | Core(neuroectoderm) | 100 |
| | Basal(mesoderm) | 15 |

| | | |
|---|---|---|
| | Basal(neuroectoderm) | 100 |
| *λ_s* | **Strength of Surface Constraint** | |
| | Strength of Apical(mesoderm) surface constraint | 1 |
| | **Apical Constriction** | |
| d_t | Target link length for apical links between apical domains in neighbouring cells | 1 pixel |
| *M_min* | Minimum strength of contractile force(myosin level) | 1 |
| M_multiplier | Peak value multiplier for contractile force(myosin level) | 15 |
| t_relax | Relaxation time after each myosin profile | 2000 MCS |
| | **Cell Stiffness Parameters** | |
| *λ_sB* | Strength of Basal-Core internal link(sub-basal stiffness) | [0,10,20,...90,100] |
| *λ_sA* | Strength of Apical-Core internal link(sub-apical stiffness) | [0,10,20,...190,200] |
| | **Contact Energies(External)** | |
| J_mm | medium-medium | 0 |
| J_ma | medium-apical | 8 |
| J_mc | medium-core | 100 |
| J_mb | medium-basal | 8 |
| J_mc' | medium-core' | 100 |
| J_mb' | medium-basal' | 8 |
| J_aa | apical-apical | 10 |
| J_ac | apical-core | 100 |
| J_ac' | apical-core' | 100 |
| J_ab | apical-basal | 100 |
| J_ab' | apical-basal' | 100 |
| J_cc' | core-core' | 10 |
| J_cc | core-core | 100 |
| J_c'c' | core'-core' | 100 |
| J_cb | core-basal | 100 |
| J_cb' | core-basal' | 100 |
| J_c'b | core'-basal | 100 |

| | | |
|---|---|---|
| J_c'b' | core'-basal' | 100 |
| J_bb' | basal-basal' | 5 |
| J_bb | basal-basal | 100 |
| J_b'b' | basal'-basal' | 100 |
| | **Contact Energies(Internal)** | |
| j_bc | basal-core | 2 |
| j_ab | apical-basal | 100 |
| j_ac | apical-core | 2 |
| j_b'c' | basal'-core' | 2 |
| j_ab' | apical-basal' | 100 |
| j_ac' | basal'-core' | 2 |
